# Metabolic engineering of stomatal precursor cells enhances photosynthetic water-use efficiency and vegetative growth under water-deficit conditions in *Arabidopsis thaliana*

**DOI:** 10.1101/2024.08.29.610053

**Authors:** Jacques W. Bouvier, Steven Kelly

## Abstract

Stomata are epidermal pores that control the exchange of gaseous CO_2_ and H_2_O between plants and their environment. Modulating stomatal density can alter this exchange, and thus presents a target for engineering improved crop productivity and climate resilience. Here, we show that stomatal density in *Arabidopsis thaliana* can be decreased by the expression of a water-forming NAD(P)H oxidase targeted to stomatal precursor cells. We demonstrate that this reduction in stomatal density occurs irrespective of whether the expressed enzyme is targeted to the cytosol, chloroplast stroma, or chloroplast intermembrane space of these cells. We reveal that this decrease in stomatal density occurs in the absence of any measurable impact on stomatal dynamics, or the efficiency or thermal sensitivity of photosynthesis. Consequently, overexpression plants exhibit a higher intrinsic water use efficiency due to an increase in CO_2_ fixed per unit water transpired. Finally, we demonstrate that this enhanced water-use efficiency translates to an improvement in vegetative growth and biomass accumulation under water-deficit conditions. Together, these results thus provide a novel approach for enhancing plant productivity through metabolic engineering of stomatal density.

## Introduction

Stomata mediate the exchange of gaseous CO_2_ and water vapour between plants and their external environment. Specifically, CO_2_ provides the primary substrate of photosynthesis and its uptake is thereby required to fuel plant growth. In contrast, loss of water vapour is integral to the transpiration stream which helps regulate internal water status, temperature, and nutrient uptake. As both CO_2_ and H_2_O gases share a single diffusion path but in opposite directions, controlling flux along this path is essential for balancing the physiological demands of the plant. Consequently, optimizing the distribution and behaviour of stomata is a central way in which plants adapt to different terrestrial habitats ^1–3^.

Given their role at the interface between plants and their environment, plants are able to modulate the diffusive conductance of stomata in response to diverse stimuli including light (quality and quantity), humidity, temperature, soil water availability, and atmospheric CO_2_ ^4^. By integrating these signals alongside natural circadian rhythms, stomata open to promote CO_2_ fixation during the day and close at night or under adverse conditions to limit excessive water loss. In addition to this complex and dynamic behaviour, stomata also exhibit remarkable developmental plasticity across species and in response to diverse environmental changes. Most notably, stomatal density inversely correlates with soil water availability ^5^, temperature ^6^, and ambient CO_2_ concentration ^7–13^, and positively correlates with relative air humidity ^14,15^ over both developmental or geological timescales. These environmentally-induced changes in density are generally linked with secondary adjustments in stomatal size (such that higher densities of stomata are generally smaller and *vice versa* ^16–19^), though density always takes precedence over size as the predominant anatomical constraint on leaf gaseous conductance ^11,20^. Thus, both dynamic and developmental changes in stomata play important roles in the growth and environmental adaptation of plants.

The considerable plasticity of stomatal form and function, coupled with the importance of stomata in regulating plant growth and environmental interactions, have together inspired multiple attempts to alter their properties for crop improvement. Pioneering efforts of Farquhar and Richards in the 1980’s used stable carbon isotope ratios to screen for enhanced water use efficiency in wheat ^21,22^. Subsequently, targeted engineering efforts have included a range of forward genetic approaches which have manipulated the distribution of stomata ^23–28^, as well as the mechanical ^29^, transport ^30,31,40–42,32–39^, and metabolic ^34,43,52–56,44–51^ properties of their guard cells. Of all these approaches, altering stomatal density has received the most attention, and a vast array of mutants with both increased ^57,58,67–69,59–66^ and decreased ^17,58,72–79,61,64,65,67–71^ stomatal densities have been engineered across diverse plant and crop species.

A common theme from approaches which have altered stomatal densities has been the genetic manipulation of the expression levels of stomatal development genes. Such targets have included the basic helix-loop-helix transcription factors SPEECHLESS (SPCH) ^65^, MUTE ^67^, and FAMA ^68^ which regulate the initiation, proliferation, and differentiation stages of the stomatal lineage, respectively. Engineering success has also been achieved by altering the expression levels of other important components of stomatal development including the EPIDERMAL PATTERNING FACTOR family of signalling peptides ^17,57,76–78,58,63,69–73,75^ alongside their receptor components (TOO MANY MOUTHS ^66^ and the ERECTA protein kinases ^62^), stomatal density and distribution 1 (SDD1) and its respective interactor components ^59,61,74^, and a host of other genes involved in hormone signalling and non-canonical stomatal developmental pathways ^60,64,79^. However, despite these considerable successes, the molecular and biochemical mechanisms that link these gene networks to changes in environmental and physiological cues remain poorly understood.

Nicotinamide adenine dinucleotide (NAD^+^) presents a potential metabolic nexus that can link physiological changes in the leaf to the genetic regulation of stomatal patterning ^80–82^. This potential role arises from recent studies which have demonstrated that changes in cellular NAD^+^ status either through exogenous NAD^+^ treatment ^81^, knockdown of mitochondrial (NDT1 and NDT2) and peroxisomal (PXN1) NAD^+^ transporters ^81,82^, or knockdown of poly(ADP-ribose)polymerase 1 gene involved in NAD^+^ recycling ^81^ all produce plants with decreased stomatal densities in *Arabidopsis thaliana*. This NAD^+^-mediated alteration in stomatal patterning is thought to occur via perturbation of abscisic acid (ABA) biosynthesis and signalling, which supresses the activity of SPCH ^83^ and in turn acts to inhibit cell entry into the stomatal lineage ^5,84–87^. This connection between NAD^+^ and ABA was postulated because changes in NAD^+^ homeostasis were associated with both transcriptional changes in ABA metabolic genes and endogenous levels of this phytohormone, and because alterations in stomatal density in mutants could be rescued by inhibition of ABA biosynthesis ^81^. Intriguingly, this also aligns with the known interaction of both NAD^+^ and ABA in regulating stomatal aperture ^88,89^. Thus, as NAD^+^ is one of the cornerstones of plant metabolism, the recent discovery that it also has a role in regulating stomatal cell differentiation provides a simple and unified route through which changes in physiology or environment directly feedback on stomatal pattering.

Water-forming NAD(P)H oxidases (NOX) present a powerful genetically-encoded tool for studying the role of redox status on organismal developmental processes. Specifically, NOX are a class of enzymes which catalyse the four-electron reduction of molecular oxygen in the presence of protons and reducing equivalents NADH and nicotinamide adenine dinucleotide phosphate (NADPH) to produce water and the corresponding NAD^+^/NADP^+^ electron acceptor in the following stoichiometry [2NAD(P)H + 4H^+^ + O_2_ → 2NADP^+^ + 2H_2_O]. In nature, these enzymes are thought to function in both oxidative stress management as well as in balancing the relative levels of oxidised and reduced forms of co-factor pools ^90^. Due to this physiological role and the fact that NOX enzymes are highly catalytically efficient, they have been exploited to alter ratios of oxidised and reduced co-factor pools in both fundamental and applied science contexts ^91–98^.

In this study, we aimed to engineer stomatal density through NOX-mediated manipulation of NAD(P)^+^/NAD(P)H status in stomatal precursor cells. We show that active monomeric NOX can be targeted to different compartments in plant cells. We further show that expression of NOX across these diverse subcellular compartments of stomatal precursor cells leads to a reduction in stomatal density without any adverse effects on stomatal function or leaf-level photosynthesis. Although maximal rates of CO_2_ assimilation are not affected in transgenic plants, the intrinsic photosynthetic water-use efficiency is improved such that more CO_2_ is fixed per unit water transpired by the leaf. Accordingly, this results in enhanced growth and biomass accumulation under water-deficit conditions, whilst maintaining wild-type levels of growth under well-watered conditions. In summary, this study thus provides a novel and tractable metabolic engineering approach to manipulate stomatal patterning as a potential target to enhance crop productivity and climate resilience.

## Results

### NOX can be expressed in plant cells and targeted to multiple subcellular locations

Diverse water-forming NAD(P)H oxidases (NOX) enzymes have been characterised across bacteria ^94,95,107–116,99,117,100–106^, archaea ^90,118–123^, and eukaryotes ^124–126^. Here, we used an engineered variant of this enzyme from the bacteria *Streptococcus mutans* (*Sm*NOX) for the basis of all experiments. This variant was chosen based on several criteria. First, *Sm*NOX exhibits NAD(P)H bifunctionality ^127^ and is thus capable of oxidising co-factor pools in a range of eukaryotic cellular contexts. Second, *Sm*NOX is functional across a broad pH and temperature range with optimal activities matching physiological conditions *in planta* ^99,128^. Third, *Sm*NOX is monomeric ^99^ allowing a higher possibility for it to tolerate protein-protein fusions and N-terminal targeting peptides whilst maintaining functional activity. Finally, *Sm*NOX exhibits negligible secondary H_2_O_2_ production that may contribute to undesirable off-target effects ^127,128^.

To assess whether NOX can be produced in plants, we first tested *Sm*NOX expression using a transient protoplast expression system. For this purpose, mesophyll protoplasts were isolated from mature rosette leaves of *A. thaliana* and transformed with a genetic construct containing the full-length *Sm*NOX coding sequence translationally fused to a C-terminal GFP (Supplemental File 1, Figure S1A). Analysis of cell fluorescence using confocal microscopy confirmed that the *Sm*NOX-GFP fusion was translated in plant cells, and that the enzyme localised to the cytosol when driven in the absence of any additional targeting sequences (Figure 1A) in agreement with a predicted bioinformatic localisation (Supplemental File 1, table S1).

**Figure 1.**
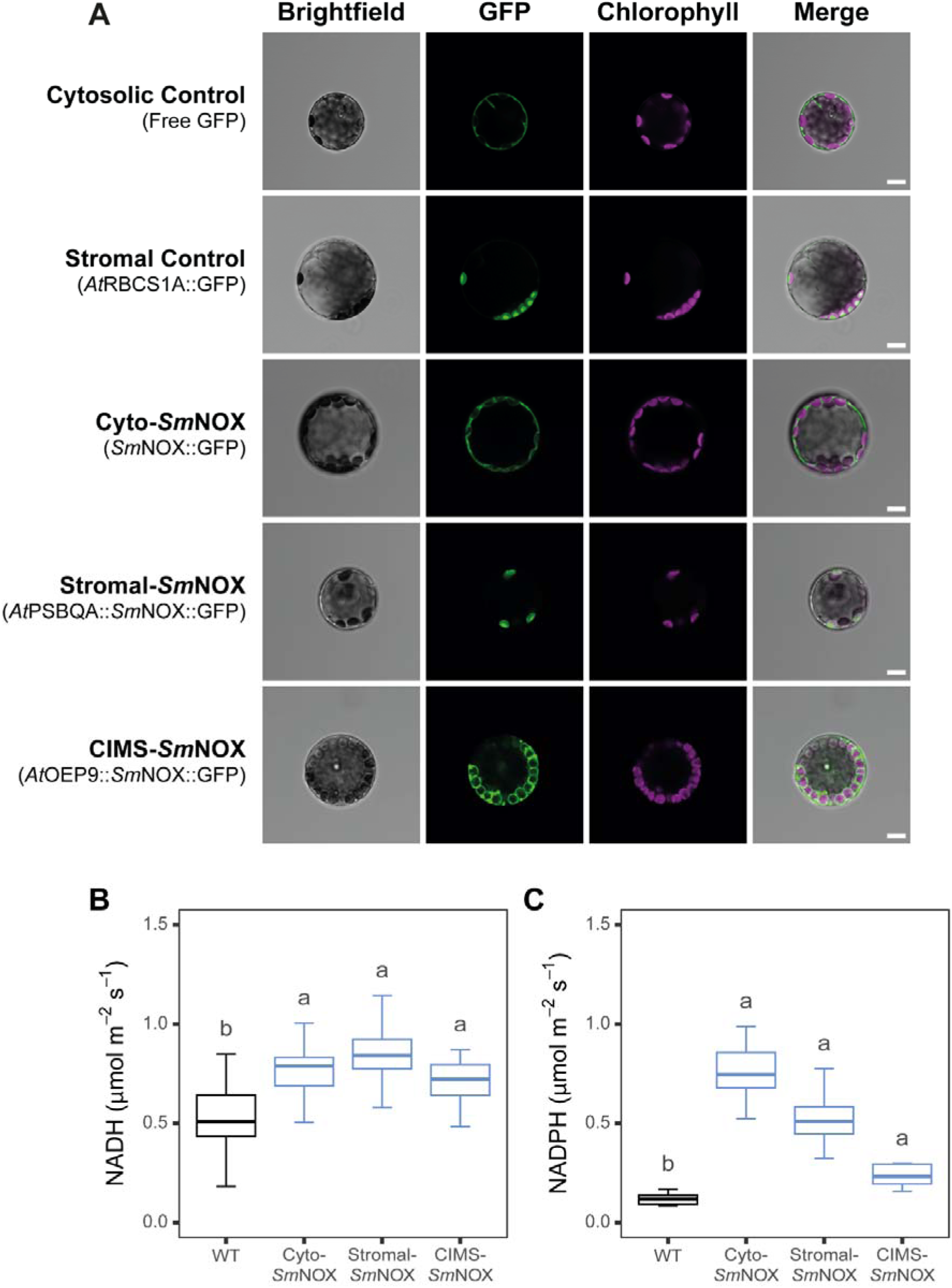
Validation of *Sm*NOX localisation and functional activity *in planta*. A) The subcellular localisation of *Sm*NOX in *Arabidopsis thaliana* protoplasts. Free GFP: protoplasts expressing eGFP (cytosolic control). *At*RBCS1A::GFP: protoplasts expressing eGFP fused to the C-terminus of the rubisco small subunit (chloroplast stroma control). Cyto-*Sm*NOX: protoplasts expressing *Sm*NOX fused at the C-terminus to eGFP (cytosolic localisation). Stromal-*SmNOX*: protoplasts expressing *SmNOX* fused at the C-terminus to eGFP and fused at the N-terminus to the chloroplast transit peptide of the photosystem II subunit QA protein (chloroplast stromal localisation). CIMS-*Sm*NOX: protoplasts expressing *SmNOX* fused at the C-terminus to eGFP and fused at the N-terminus to the full length outer envelope protein 9 (CIMS; chloroplast intermembrane space localisation). Three separate signals are shown. Green: eGFP fluorescence. Violet: Thylakoid-derived chlorophyll autofluorescence. Grey: bright-field. The merged channel represents all three signals overlaid in a single image. Scale bar = 10 µM. B) Boxplot depicting the rate of NADH oxidation from crude leaf lysate of wild-type (black) and transgenic plants (blue). Abbreviations follow that described in (A). Data represent an average across three independent single copy lines (*n* = 4 plants per line). C) As in (B), but for the total NADPH oxidation from crude leaf lysate (*n* = 5 plants per line). Differences between transgenic plants and WT are assessed by Fisher LSD post-hoc analysis following a two-way ANOVA, where letters above each box represent statistically significant differences in mean values (p ≤ 0.05). The raw data can be found in Supplemental File 4.

Whilst NAD^+^ is produced in the cytosol, the regulatory mechanism that governs cytosolic NAD^+^ levels is dependent on chloroplast NADP^+^ production. Accordingly, we sought to test whether *Sm*NOX could also be localised closer to the site of NADP^+^ supply in both the chloroplast stroma (the site of NADPH production) and the chloroplast intermembrane space (the site of NADPH/NADH transfer via the malate valve). *Sm*NOX was successfully localised to the chloroplast stroma (stromal-*Sm*NOX) when fused to the transit peptide from the photosystem II subunit QA (PSBQA) protein (Figure 1A). This stromal localisation was verified by comparison to a stromal control consisting of the full-length rubisco small subunit gene fused to GFP (Figure 1A). Moreover, *Sm*NOX was also successfully targeted to the chloroplast intermembrane space (CIMS-*Sm*NOX) when fused to the full-length outer envelope protein 9 gene (Figure 1A). In summary, *Sm*NOX can be expressed in plants and can be localised to diverse subcellular compartments including the cytosol (the site of NAD^+^ production), chloroplast stroma (the site of NADP^+^ production), and chloroplast intermembrane space (the site of NAD^+^/NADP^+^ transfer).

### Identifying a promoter capable of driving stable and high expression of SmNOX in stomatal precursor cells

To investigate whether *Sm*NOX could affect plant stomatal patterning, it was necessary to next find a promoter which drives high gene expression in undifferentiated stomatal progenitor cells in the leaf epidermis (i.e., during the developmental window prior to onset of stomatal lineage differentiation) (Supplemental File 1, Figure S2A). For this purpose, we analysed a cell-specific transcriptome dataset derived from all major cell types spanning the stomatal developmental trajectory ^129^. This revealed that the expression of genes encoding known master-regulators of stomatal development including the transcription factors SPEECHLESS (SPCH), MUTE, and FAMA were expressed at low levels which were considered to be insufficient to achieve *Sm*NOX-mediated alteration of cellular NAD^+^ status (Supplemental File 1, Figure S2B – S2J). Consequently, we sought to identify an alternative more highly expressed promoter.

Interrogation of the above transcriptome data revealed that the gene encoding the chlorophyll a/b binding protein 3 (CAB3) exhibited high mRNA abundance in the target epidermal tissue of interest (Supplemental File 1, Figure S2K). Specifically, CAB3 was ranked as the 36^th^ (out of ∼28,000) most highly expressed nuclear-encoded gene in this cell type (Supplemental File 1, Figure S2L and S2M), and exhibited transcripts levels that were ∼1,800-fold more abundant than SPCH (0.5 TPM), ∼9,000-fold more abundant than MUTE (0.1 TPM), and ∼425-fold more abundant than FAMA (2.1 TPM), respectively (Supplemental File 1, Figure S2B – S2M). Thus, as CAB3 is one of the most abundant transcripts in the cell type of interest, and as CAB3 is a well-characterised promoter that is extensively used for driving transgene expression in plant synthetic biology ^130^, this promoter was selected for use herein. Genetic constructs containing targeted and untargeted variants of the *Sm*NOX coding sequence were thus cloned under the control of the CAB3 promoter and were transformed into *A. thaliana* ecotype Col-0 (Supplemental File 1, Figure S1B and Figure S3A and S3B). Homozygous plants from three single-copy insertion events of cyto-SmNOX, stromal-SmNOX and CIMS-SmNOX were used for the basis of all subsequent experiments.

### Validation of SmNOX expression and function

To assess the functional activity of NOX in plants, total rates of NADH and NADPH oxidation were determined using *in vitro* assays containing crude lysate from mature rosette leaves. As plants possess multiple native enzymes capable of oxidising NADH and NADPH, background oxidation of NADH and NADPH was observed in all cases as expected (Figure 1B and 1C). However, the rates of both NADH and NADPH oxidation were significantly increased across all transgenic lines compared to wild-type controls (Figure 1B and 1C). Thus, there was successful expression of active *Sm*NOX in each of the targeted subcellular localisations.

It should be noted that the CAB3 promoter also drives high levels of transgene expression in other photosynthetic cells of the mature leaf (Supplemental File 1, Figure S2K – S2M and Figure S4A – S4C). Thus, although our target tissue for expression of *Sm*NOX is the leaf epidermis, only a portion of the above enzyme activity in the crude leaf lysate corresponds to this cell layer. Moreover, the rates of *Sm*NOX-mediated NADH/NADPH oxidation measured above (∼ 1.0 μmol m^−2^ s^−1^) are substantial in the context of an epidermal cell, they are not expected to impact NADH/NADPH pools in photosynthetic mesophyll cells. This is because of both the large pool size of NADH (24 – 55 μM) and NADPH (140 – 180 μM) ^131^ in the mesophyll tissue, as well as the extraordinary high rates of photosynthetic NADPH production (∼70 μmol g^−1^ s^−1^) ^132^ which occurs during the light. Thus, we hypothesized that *Sm*NOX would be capable of manipulating endogenous levels of NAD^+^ in leaf cells with low photosynthetic activity such as epidermal cells, but would have little or no effect on the NAD^+^ status or photosynthesis of mesophyll cells.

### SmNOX activity causes reductions in stomatal density

Given that *Sm*NOX was active in the transgenic plants, we next investigated whether this resulted in an alteration in stomatal patterning. For this purpose, the epidermal composition of the abaxial surface of mature rosette leaves from 8-week-old plants was analysed by bright-field microscopy. This analysis revealed that stomatal densities in all *Sm*NOX-expressing lines were reduced by 20 – 30% compared to wild-type control plants (Figure 2A). Thus, changes in cellular NAD(P)^+^/NAD(P)H ratios, irrespective of the location of perturbation of that ratio, caused a suppression in stomatal development resulting in a reduced density of stomata on leaves.

**Figure 2.**
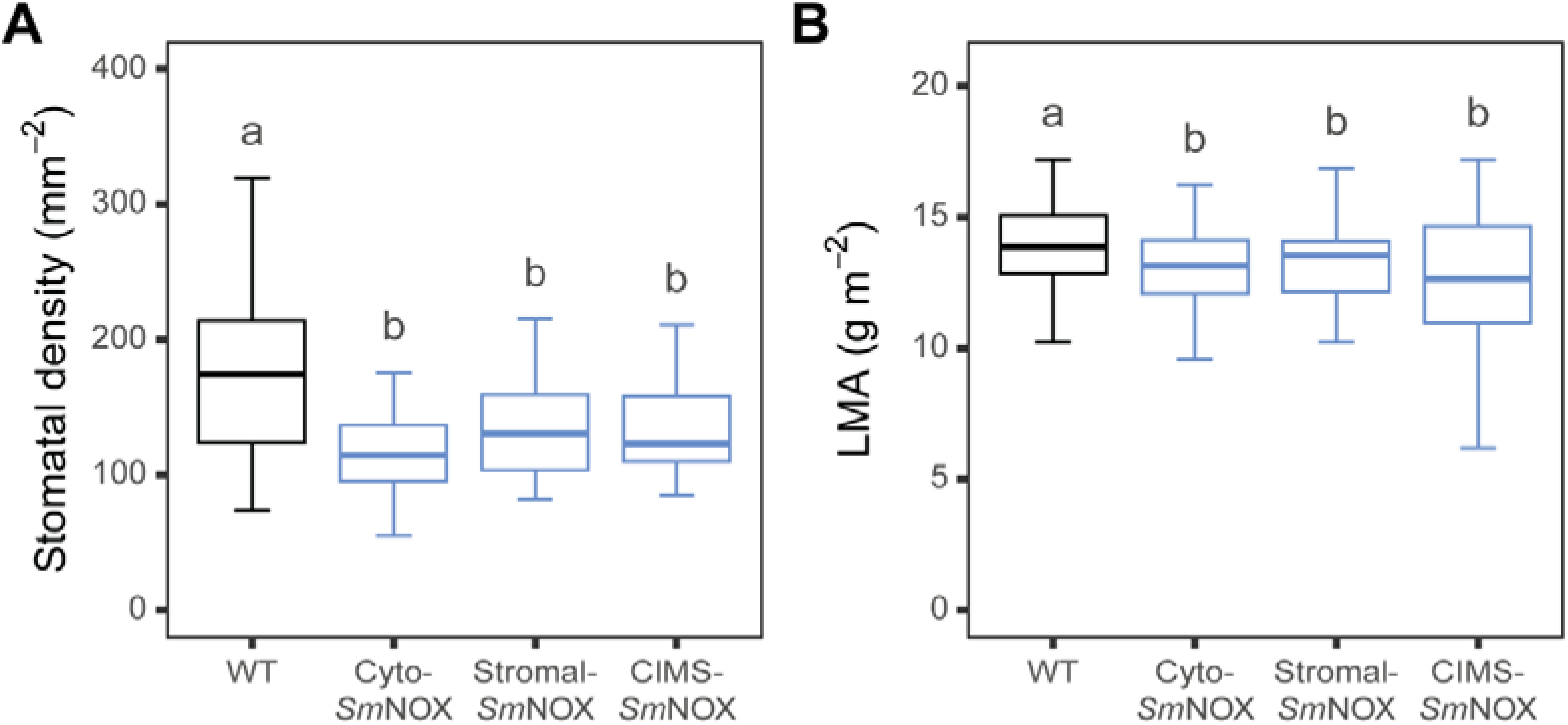
The effect of *Sm*NOX expression on stomatal density. A) Boxplot depicting the stomatal density (count mm^−2^) measured on the abaxial surface of mature rosette leaves from wild-type (black) and transgenic plants (blue). WT: wild-type. Cyto-*Sm*NOX: cytosolic *Sm*NOX. Stromal-*Sm*NOX: chloroplast stroma *Sm*NOX. CIMS-*Sm*NOX: chloroplast intermembrane space *Sm*NOX. Data represent an average across three independent single copy lines (*n* = 9 – 10 plants per line). B) As in (A), but for the measured leaf mass per area (LMA, g dry mass per m^−2^ leaf area) of mature rosette leaves (*n* = 10 plants per line). Differences between transgenic plants and WT are assessed by Fisher LSD post-hoc analysis following a two-way ANOVA, where letters above each box represent statistically significant differences in mean values (p ≤ 0.05). The raw data can be found in Supplemental File 4.

Stomatal density changes have previously been linked with secondary changes in mesophyll development (such that leaves with reduced stomatal numbers are generally thinner and *vice versa* ^133,134^) which can have important consequences for plant physiology ^135–141^. Thus, we also investigated whether *Sm*NOX-expression was associated with an alteration in leaf width or cellular density by measuring leaf mass per area (LMA = leaf dry mass / leaf area). This revealed *Sm*NOX-expressing lines exhibited a small but significant reduction in LMA compared to wild-type (Figure 2B). Thus, expression of *Sm*NOX results in both a decrease in stomatal patterning and anticipated secondary alterations in leaf architecture.

### SmNOX-induced changes in stomatal density improve plant photosynthetic water-use efficiency

As stomatal density is a key determinant of leaf gaseous conductance, we next characterised the effect of *Sm*NOX expression on plant physiological performance. To test this, we first assessed the photosynthetic parameters of *Sm*NOX-expressing lines along a light intensity gradient using an infrared open-gas exchange system (Figure 3A – 3C). This analysis revealed that there was a small reduction in CO_2_ assimilation rate (*A*) between *Sm*NOX-expressing lines and wild-type plants (Figure 3A). However, stomatal conductance (*g*_s_) was substantially reduced across all *Sm*NOX-expressing lines (Figure 3B), consistent with these mutants exhibiting lower ratios of intercellular CO_2_ to ambient CO_2_ (*C*_i_/*C*_a_) and reduced rates of transpiratory water loss under high light (Supplemental File 1, Figure S5A and S5B). *Sm*NOX-expressing lines also exhibited comparable light-saturated rates of CO_2_ assimilation (*A*_sat_, Figure 3D, Supplemental File 1, table S2), and substantially lower light-saturated rates of stomatal conductance (*g*_s sat_, Figure 3E, Supplemental File 1, table S2). As a consequence of these combined effects on *A* and *g*_s_, all *Sm*NOX-expressing lines exhibited a higher intrinsic water use efficiency (*iWUE = A*/*g_s_*) across a broad range of light intensities (Figure 3C and 3F, Supplemental File 1, table S2).

**Figure 3.**
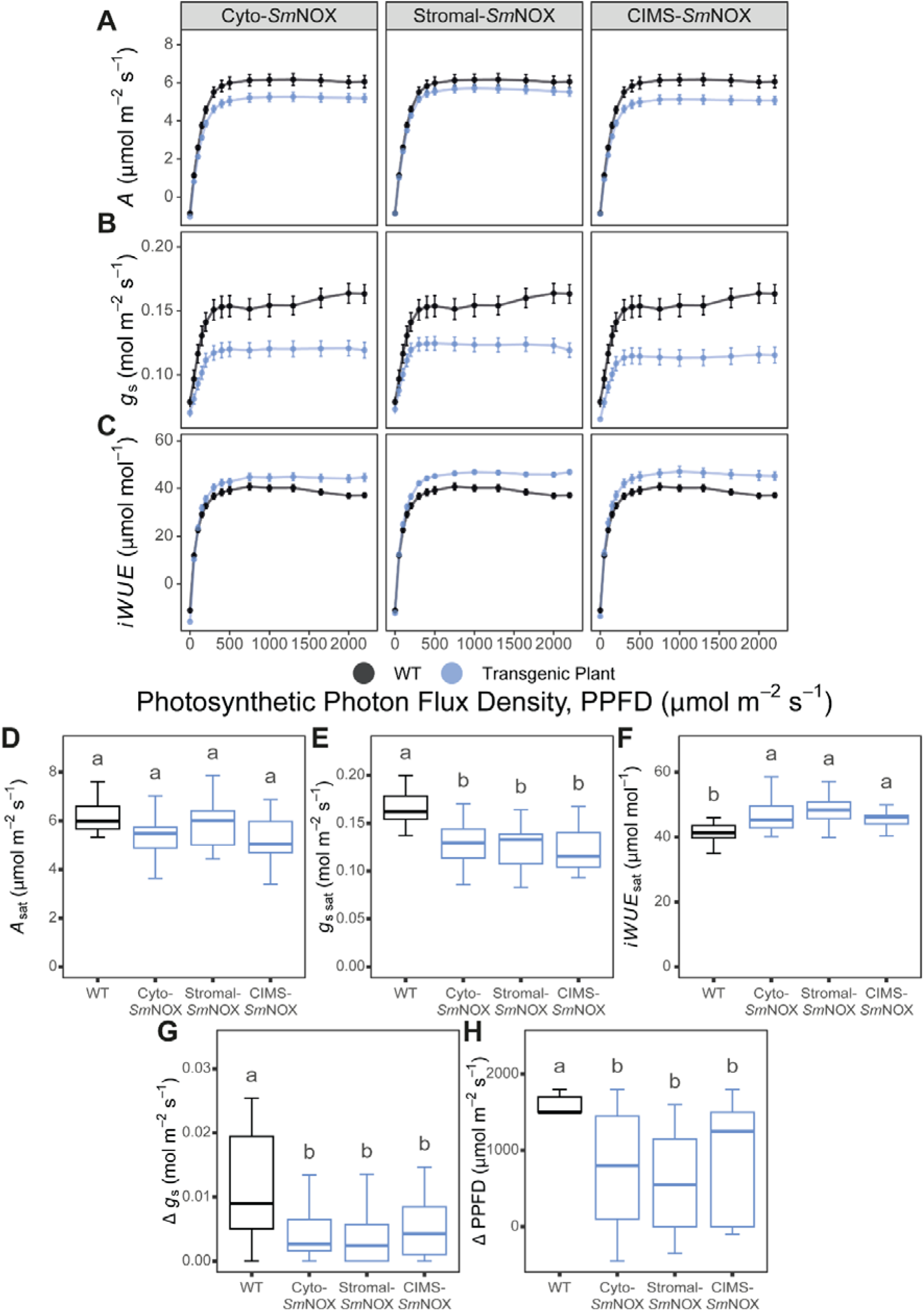
The response of plant gas exchange to light. A) CO_2_ assimilation rate (*A*, μmol m^−2^ s^−1^) at different PPFD in wild-type (black) and transgenic plants (blue). WT: wild-type. Cyto-*Sm*NOX: cytosolic *Sm*NOX. Stromal-*Sm*NOX: chloroplast stromal *Sm*NOX. CIMS-*Sm*NOX: chloroplast intermembrane space *Sm*NOX. Data represent mean ± 1 S.E. B) as in (A), but for stomatal conductance (*g*_s_, mol m^−2^ s^−1^). C) as in (A), but for intrinsic water-use efficiency (*iWUE*, μmol mol^−1^). D) Boxplot depicting the light-saturated assimilation rate (*A*_sat_, μmol m^−2^ s^−1^). The colour scheme and abbreviations follow that described in (A). E) As in (D), but for the light-saturated stomatal conductance (*g*_s sat_, mol m^−2^ s^−1^). F) As in (D), but for the light-saturated intrinsic water-use efficiency (*iWUE*_sat_, μmol mol^−1^). G) Boxplot depicting the difference in stomatal conductance between the light intensity at which 95% maximum assimilation rate and the light-saturated stomatal conductance are achieved (Δ *g*_s_, mol m^−2^ s^−1^). H) As in (G), but for the difference in light intensity at which 95% maximum assimilation rate and the light-saturated stomatal conductance are achieved (Δ PPFD, μmol m^−2^ s^−1^). All data represent an average across three independent single copy lines (*n* = 7 – 8 plants line). Differences between transgenic plants and WT are assessed by Fisher LSD post-hoc analysis following a two-way ANOVA, where letters above each box represent statistically significant differences in mean values (p ≤ 0.05). The raw data can be found in Supplemental File 4.

In addition to an enhanced *iWUE*, *Sm*NOX expression also resulted in tighter coordination between rates of *A* and *g*_s_. Specifically, all mutants displayed a reduced “overshooting” of *g*_s_ at light intensities above those where 95% maximum assimilation was achieved (beyond which neither CO_2_ delivery nor light is considered limiting) (Supplemental File 1; Figure 3G and 3H; Supplemental File 1, Figure S6A – S6F, table S3). This improved *iWUE* and tighter coupling between *A* and *g*_s_ was not associated with a change in any other measured photosynthetic parameter including light compensation point, mitochondrial respiration in the light, apparent quantum yield (i.e., the efficiency by which light can be converted into ATP and NADPH), or the light-saturated rate of electron transport (Supplemental File 1, Figure S5C – S5F, table S2). Moreover, there was no consistent difference in any derived photosynthetic parameter when CO_2_ assimilation was measured as a function of sub-stomatal CO_2_ concentration (Supplemental File 1, Figure S7, Table S4). This included no change in the maximal CO_2_ saturated assimilation rate, CO_2_-compensation point, the transition point between rubisco-limited and RuBP regeneration-limited rates of photosynthesis, the rubisco Michaelis constant for CO_2_ in the presence of 21% O_2_ air, the carboxylation rate of rubisco, or the maximum electron transport rate through PSII (Supplemental File 1, Figure S6, Table S4). As anticipated, NOX activity was able to alter stomatal density, but was not sufficient to exert any measurable impact on mesophyll cell photosynthesis. Accordingly, *Sm*NOX-mediated reduction in stomatal density drives both an enhanced photosynthetic water-use efficiency and a tighter coordination between rates of CO_2_ assimilation and gas exchange.

### SmNOX does not affect the kinetics of stomatal movement

We next sought to examine whether changes in stomatal density caused by *Sm*NOX expression were also associated with off-target impacts on stomatal behaviour. Specifically, we chose to assess the speed and magnitude of stomatal responses to light, given that this is one of the most important fluctuating variables that plants face under real-world conditions in the field. Light was also chosen because NADP^+^ is the terminal electron acceptor in the photosynthetic electron transport chain and thus the impact of light on stomatal kinetics had potential to be perturbed by *Sm*NOX expression. For the purpose of this analysis, the experimental design of ^142^ was followed. In brief, plants were first acclimated at 100 μmolLJm^−2^LJs^−1^ of light for 60 minutes, followed by a step increase in light intensity to 1000 μmolLJm^−2^LJs^−1^ for 60 minutes, and a final step decrease in light intensity to 100 μmolLJm^−2^LJs^−1^ for 30 minutes, with gas-exchange parameters recorded at one-minute intervals over this entire period.

This analysis revealed that all plants experience a normal sigmoidal response of *g*_s_ in response to step changes in irradiance (Figure 4A and 4B, Supplemental File 1, Figure S8). Moreover, subsequent model-based interrogation ^143^ of these data demonstrated that neither the amplitude nor the speed of stomatal responses was different between wild-type and transgenic plants. Specifically, no difference was found in either the modelled initial lag time in *g*_s_, the maximum rate of change of *g*_s_ (calculated from the slope of the exponential phase of the curve), or the time taken for *g*_s_ to reach new steady-state upon the initial step increase in light intensity from 100 to 1000 μmolLJm^−2^LJs^−1^ (Figure 4C – 4E, Supplemental File 1, table S5). Similarly, no difference could be detected in the time taken to reach new steady-state *g*_s_ upon the subsequent step decrease in light intensity from 1000 to 100 μmolLJm^−2^LJs^−1^ (Figure 4F, Supplemental File 1, table S5). Thus, consistent with the lack of effect of *Sm*NOX on plant photosynthesis, *Sm*NOX expression did not impair the dynamic behaviour of stomatal movement.

**Figure 4.**
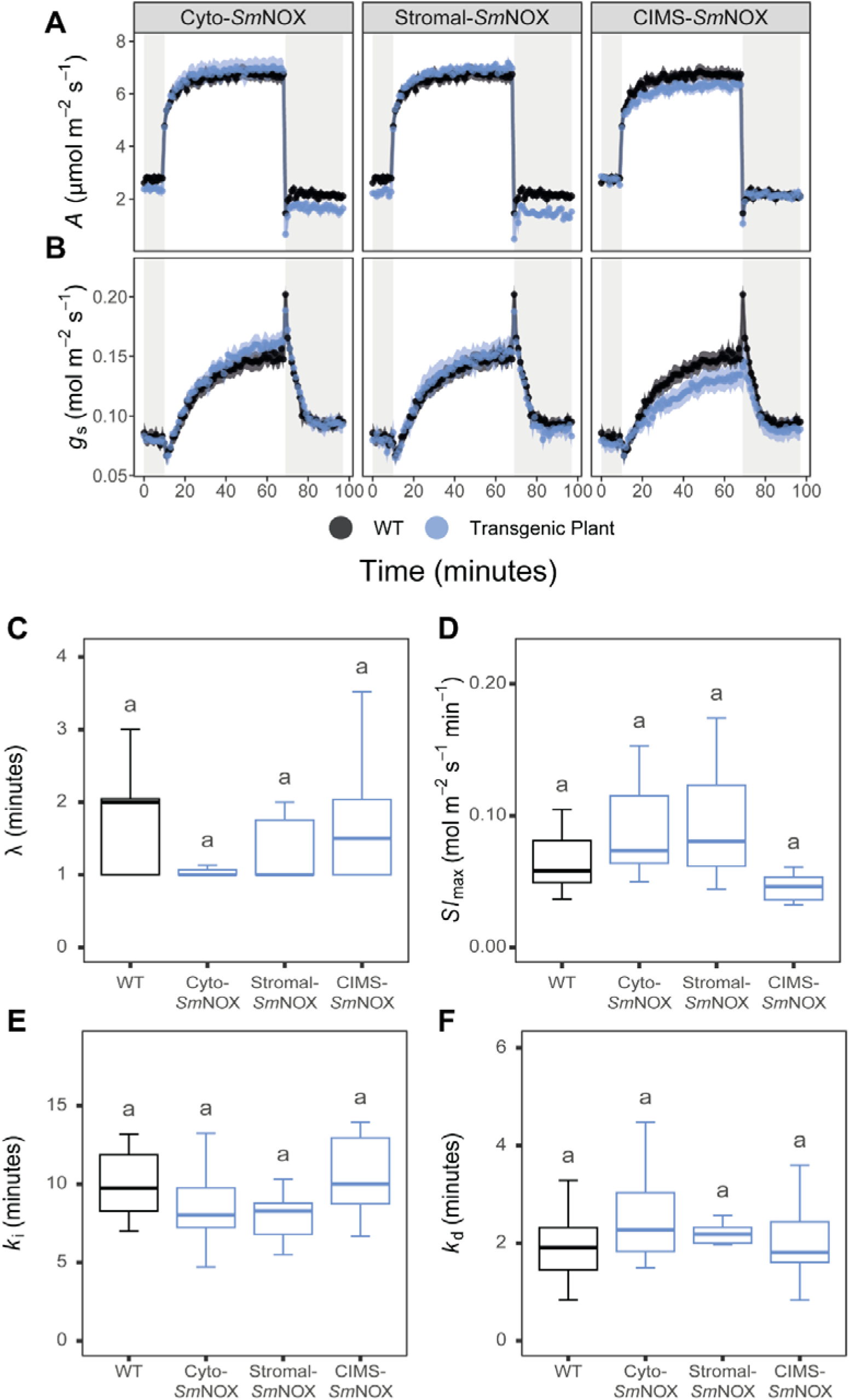
The temporal response of plant gas exchange to step changes in light intensity. A) The response of CO_2_ assimilation rate (*A*, μmol m^−2^ s^−1^) to step changes in light intensity in wild-type (black) and transgenic plants (blue). Shaded and unshaded areas represent periods of light intensity of 100 μmol m^−2^ s^−1^ and 1000 μmol m^−2^ s^−1^, respectively. WT: wild-type. Cyto-*Sm*NOX: cytosolic *Sm*NOX. Stromal-*Sm*NOX: chloroplast stromal *Sm*NOX. CIMS-*Sm*NOX: chloroplast intermembrane space *Sm*NOX. Data represent mean ± 1 S.E. B) as in (A), but for stomatal conductance (*g*_s_, mol m^−2^ s^−1^). C) Boxplot depicting the estimated initial lag in response time of *g*_s_ (λ, minutes) after the step increase in light intensity from 100 μmol m^−2^ s^−1^ to 1000 μmol m^−2^ s^−1^. The colour scheme and abbreviations follow that described in (A). D) As in (C), but for the estimated maximal rate of stomatal opening (*Sl* _max_, mol m^−2^ s^−1^ min^−1^) after the step increase in light intensity from 100 μmol m^−2^ s^−1^ to 1000 μmol m^−2^ s^−1^. E) As in (C), but for the estimated time taken to achieve new steady state stomatal conductance (*k*_i_, minutes) after the step increase in light intensity from 100 μmol m^−2^ s^−1^ to 1000 μmol m^−2^ s^−1^. F) As in (C), but for the estimated time taken to achieve new steady state stomatal conductance (*k*_d_, minutes) after the step decrease in light intensity from 1000 μmol m^−2^ s^−1^ to 100 μmol m^−2^ s^−1^. All data represent an average across three independent single copy lines (*n* = 3 – 4 plants per line). Differences between transgenic plants and WT are assessed by Fisher LSD post-hoc analysis following a two-way ANOVA, where letters above each box represent statistically significant differences in mean values (p ≤ 0.05). The raw data can be found in Supplemental File 4.

### SmNOX does not alter plant susceptibility to supra-optimal temperatures

One important function of the plant transpiration stream is to mediate the temperature control of plant aerial tissues through evaporative cooling. As such, plants exhibiting reduced stomatal numbers might be more susceptible to thermal damage of photosynthesis under supra-optimal temperatures, owing to impairment of their temperature control mechanism (Supplemental File 1, Figure S5B). To investigate this, we measured plant photosynthesis across step-wise increases in air temperatures from 20 to 45 °C. Under these conditions no difference could be detected in any photosynthesis parameter at any temperature between *Sm*NOX-expressing lines and wild type plants. In all plants, the rate of *A* increased with temperature from 20 °C, reaching an optima around 30 °C, and then rapidly declining with increasing temperatures beyond this optima consistent with thermal-induced inactivation of the photosynthetic machinery (Figure 5A). Despite an attenuated rate of *g*_s_ in mutant plants across all conditions, a comparable trend was also observed in this between both mutant and wild-type plants (Figure 5B). Moreover, no difference could be detected in the measured leaf temperature of *Sm*NOX-expressing lines compared to wild-type plants at any given air temperature (Figure 5C). Thus, reductions in stomatal density in *Sm*NOX-expressing lines was not associated with either a change in leaf temperature or any change in the susceptibility to thermal damage of the photosynthetic machinery under supra-optimal temperatures.

**Figure 5.**
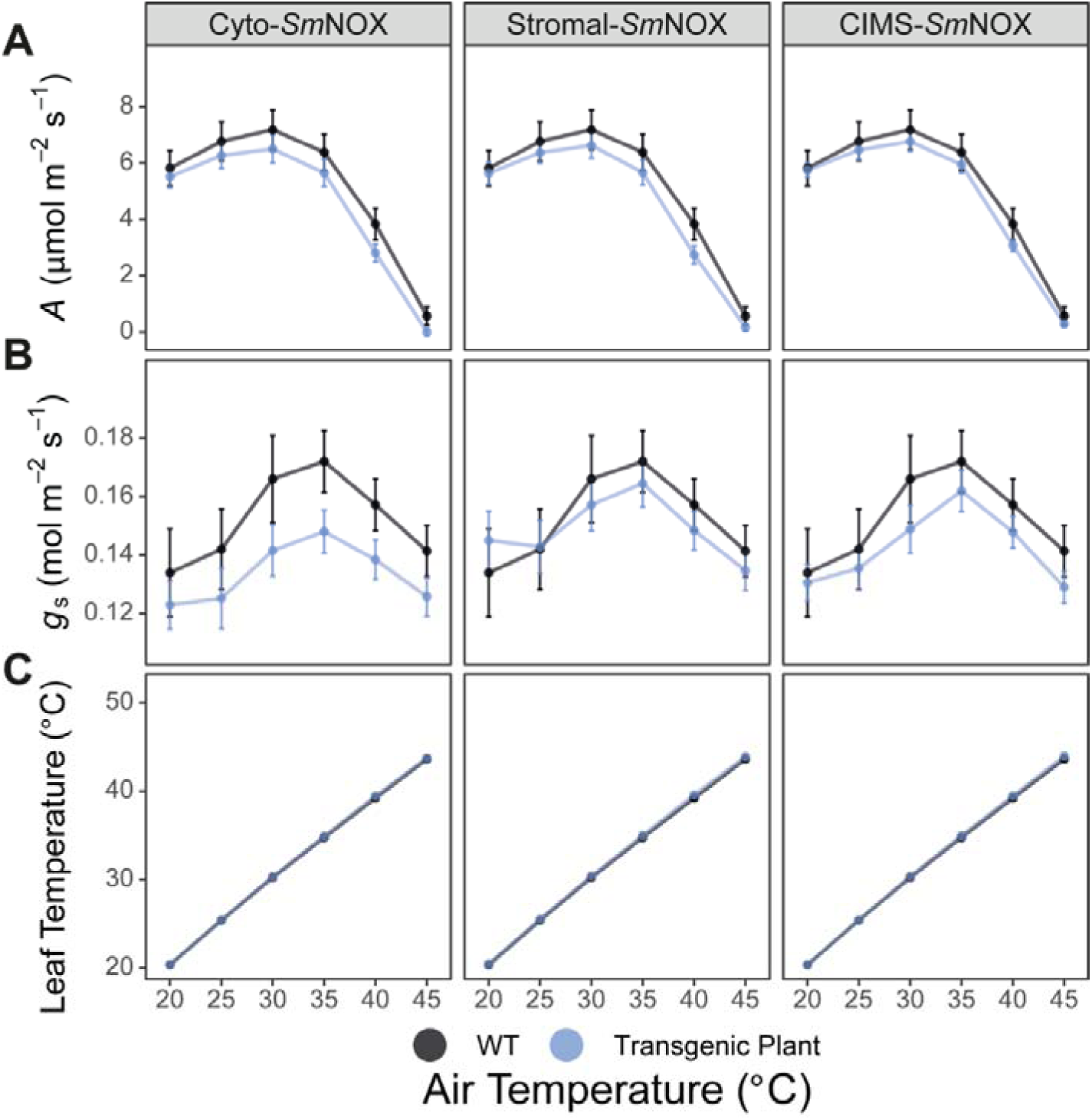
The response of plant gas exchange to air temperature (°C). A) CO_2_ assimilation rate (*A*, μmol m^−2^ s^−1^) at different air temperatures in wild-type (black) and transgenic plants (blue). WT: wild-type. Cyto-*Sm*NOX: cytosolic *Sm*NOX. Stromal-*Sm*NOX: chloroplast stromal *Sm*NOX. CIMS-*Sm*NOX: chloroplast intermembrane space *Sm*NOX. Data represent mean ± 1 S.E. B) as in (A), but for stomatal conductance (*g*_s_, mol m^−2^ s^−1^). C) as in (A), but for leaf temperature (°C). All data represent an average across three independent single copy lines (*n* = 3 – 5 plants per line). The raw data can be found in Supplemental File 4.

### SmNOX improves vegetative growth and biomass accumulation under water-deficit conditions

Based on the findings that *Sm*NOX expression caused an enhancement in *iWUE* without any off-target effects on photosynthetic efficiency, stomatal behaviour, or thermal sensitivity, we next sought to determine whether an impact on plant growth could also be observed. For this purpose, we grew plants in a fully randomised block design under long-day conditions with a 16-hour photoperiods and 250 μmol m^−2^ s^−1^ non-limiting light. Under these conditions, no difference in the vegetative growth or performance between transgenic *Sm*NOX-expressing plants and wild-type plants was detected under well-watered conditions (Supplemental File 1, Figure S9A – S9C). Given the higher *iWUE* of mutant plants, we next determined whether these improve photosynthesis-water relations resulted in any change in plant growth under water-deficit conditions. For this purpose, plant trays were watered prior to stratification, but water was then withheld for 15-days after the termination of stratification to maintain sub-optimal and deteriorating soil-water saturation levels during vegetative growth (after which watering was resumed when the top layer of soil was visibly dry every ∼7 days). This assay revealed that *Sm*NOX expressing plants produced more biomass and larger rosettes compared to wild-type plants over equivalent periods of time (Figure 6A-E). This result was caused by *Sm*NOX expressing-lines maintaining higher relative growth rates during water-deficit (Figure 6F). Consistent with this enhancement in vegetative growth and tolerance to adverse water conditions, mutants also exhibited earlier times of bolting (Figure 6G) and flowering (Figure 6H), and produced visibly taller inflorescences relative to wild-type (Figure 6I). However, these alterations in vegetative growth and biomass did not translate to improved yields, and all *Sm*NOX expressing-lines exhibited a reduction in total seed mass (Figure 6J). Thus, *Sm*NOX-expressing plants exhibited enhanced biomass production and tolerance to water-deficit conditions but an overall impairment of seed production.

**Figure 6.**
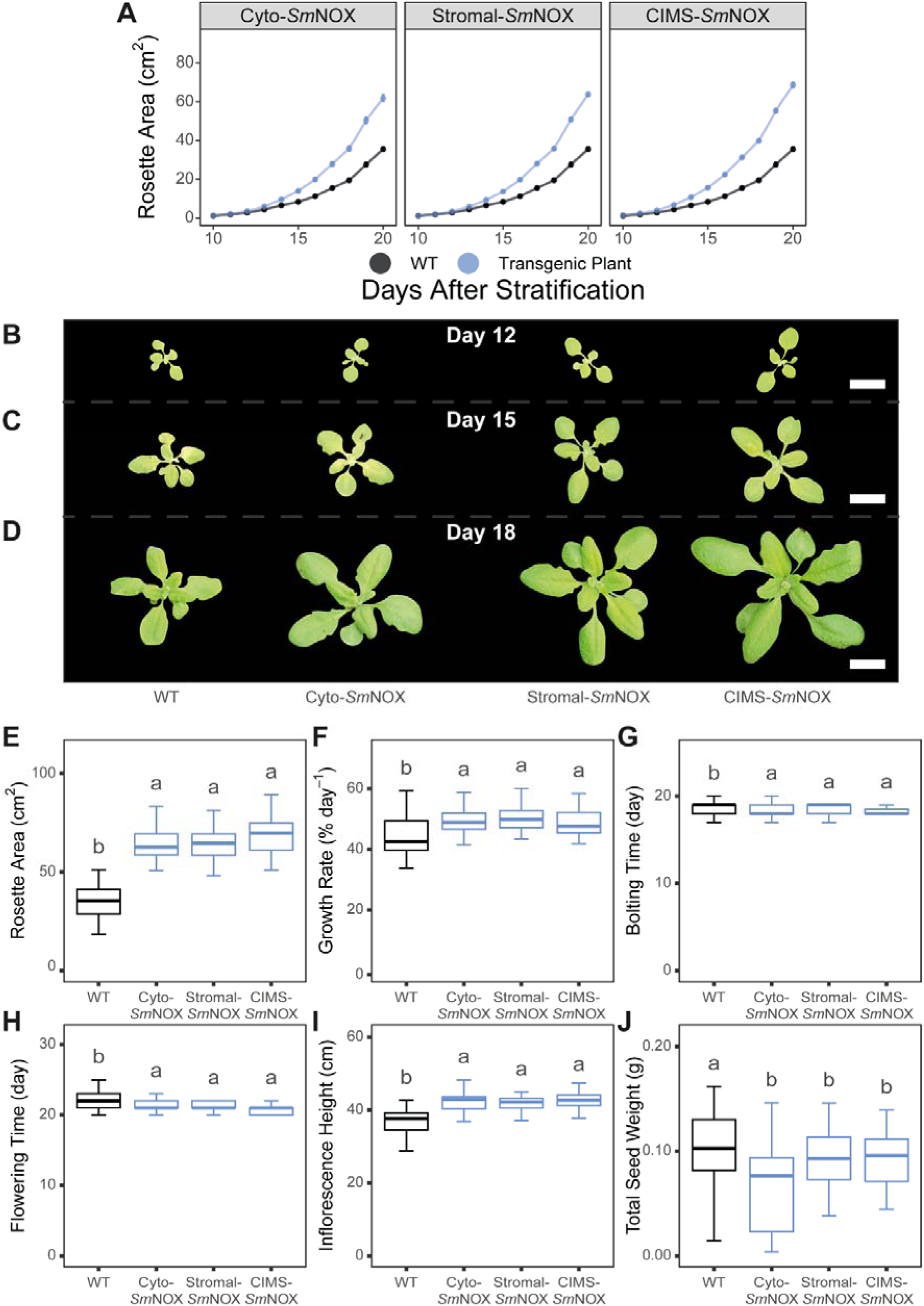
The vegetative and reproductive performance of plants under water-deficit conditions. (A) The increase in visible rosette area (cm^2^) over time in wild-type (black) and transgenic plants (blue). WT: wild-type. Cyto-*Sm*NOX: cytosolic *Sm*NOX. Stromal-*Sm*NOX: chloroplast stromal *Sm*NOX. CIMS-*Sm*NOX: chloroplast intermembrane space *Sm*NOX. Data represent mean ± 1 S.E. B) Representative images of wild-type and transgenic plants taken on day 12 post stratification. Scale bar = 2.5 cm. C) As in (B), but for day 15 post stratification. D) As in (B), but for day 18 post stratification. In parts B – D, the same individual plant is shown across the growing period and all images were generated by computationally removing background from the aerial photographs analysed for growth analyses. E) Boxplot depicting the maximum visible rosette area at the end of vegetative growth (day = 20). The colour scheme and abbreviations follow that described in (A). F) As in (E), but for the average percentage growth rate of plants computed over the vegetative growth phase (% increase in rosette area day^−1^). G) As in (E), but for the time taken for bolting to occur in plants (days post stratification). H) As in (E), but for the time taken for flowering to occur in plants (days post stratification). I) As in (E), but for the maximum inflorescence height (cm) measured at day 32 post stratification. J) As in (E), but for the total seed weight harvested from plants after drying (g). All data represent an average across three independent single copy lines (*n* = 17 – 20 plants per line for all measurements, except total seed weight where *n* = 11 – 13 plants per line). Differences between transgenic plants and WT are assessed by Fisher LSD post-hoc analysis following a two-way ANOVA, where letters above each box represent statistically significant differences in mean values (p ≤ 0.05). The raw data can be found in Supplemental File 4.

## Discussion

Manipulation of stomatal density is a key target for enhancing crop productivity and climate resilience under current and future conditions ^144,145^. In the present study, we demonstrate that driving the expression of a water-forming NAD(P)H oxidase from *Streptococcus mutans* (*Sm*NOX) in stomatal precursor cells causes a reduction in stomatal density in *Arabidopsis thaliana*. We show that this reduced stomatal density phenotype occurs irrespective of whether *Sm*NOX is targeted to the cytosol, chloroplast stroma, or chloroplast intermembrane space. We also find that neither *Sm*NOX expression nor *Sm*NOX-mediated reductions in stomatal density had any detectable impact on plant CO_2_ assimilation rate, the efficiency or thermal sensitivity of photosynthesis, the kinetics of stomatal movement, or plant growth under well-watered conditions. Nonetheless, we discover that all *Sm*NOX-expressing lines exhibit an enhanced intrinsic water-use efficiency. We also show that this improvement in water-use efficiency translates to an increase in plant vegetative growth and biomass accumulation under water-deficit conditions. Collectively, we thus provide a novel metabolic engineering approach which can be leveraged to alter stomatal patterning thus plant photosynthesis-water relations.

Nicotinamide adenine dinucleotide [NAD(H)] and nicotinamide adenine dinucleotide phosphate [NADP(H)] are ubiquitous coenzymes that facilitate cellular energy transfer, redox homeostasis, and signalling in living organisms ^146^. Alongside these diverse roles, NAD^+^ has recently been implicated as a negative regulator of stomatal development. Evidence in support of this hypothesis has arisen because perturbations in cellular redox status associated with translational repression of mitochondrial (*At*NDT1 and *At*NDT2) and peroxisomal (*At*PXN) NAD^+^ transporter proteins, as well as poly (ADP-ribose) polymerase (PARP) proteins (PARP1, PARP2 and PARP3) involved in NAD^+^ recycling, have all produced *A. thaliana* mutant plants with reduced stomatal densities ^81,82^. The mechanism of this developmental effect is proposed to occur via abscisic acid (ABA) which is known to inhibit entry into the stomatal lineage by suppressing the activity of SPEECHLESS (SPCH) ^84^, possibly via induction of the cyclin-dependent kinase (CDK) inhibitor gene ICK1 ^87,147^. This interaction between NAD^+^ and ABA was postulated because NAD^+^ transport and metabolism mutants were associated with both transcriptional changes in ABA metabolic genes and reduced SPCH expression ^81^. Importantly, exogenous NAD^+^ applications were also sufficient to drive a reduction in stomatal patterning in wild-type *Arabidopsis*, but not in mutants impaired in ABA biosynthesis or signalling ^81^. In the present study, we build on this foundational biology by demonstrating that expression of a water-forming NAD(P)H oxidase in stomatal precursor cells is capable of driving a reduction in stomatal density through perturbation of endogenous cellular redox status. We further show that such a reduction in stomatal density is achieved irrespective of the subcellular compartment in which this enzyme is targeted. Thus, these results collectively demonstrate that NAD^+^ is a viable metabolic handle which can be turned to alter stomatal patterning in synthetic biology applications.

The density of stomata imposes an upper anatomical constraint on leaf diffusive conductance and thus has important physiological implications for plant productivity ^148^. Despite reduced stomatal density mutants having been generated in a diverse number of disparate species and studies ^17,58,72–79,81,82,61,64,65,67–71^, the effects of this developmental alteration on plant photosynthesis, water-use efficiency, growth, and yield traits have proven variable across different studies and plant species ^149^. In the present study, we characterise the gas exchange properties of our reduced stomatal density mutants across a range of environmental conditions to shed light on their underlying physiology. We demonstrate that these reduced stomatal density plants exhibit an increased intrinsic water-use efficiency (*iWUE*) by maintaining wild-type rates of CO_2_ assimilation at lower respective stomatal gaseous conductance. Reductions in stomatal density also caused a tighter co-ordination between the rates of CO_2_ assimilation and stomatal conductance, such that a reduced extent of water loss occurred after light-saturated rates of CO_2_ assimilation were achieved. However, decreasing stomatal density did not result in any change in the efficiency or thermal sensitivity of photosynthesis and CO_2_ assimilation, or the kinetics of stomatal movements.

Together, these results align with recent findings from Epidermal Patterning Factor mutants which similarly report reductions in stomatal density without any impact on plant CO_2_ assimilation ^58,71,72,150^. The biological basis of these collective observations is intriguing. As the rate of photosynthesis at any one time is constrained by the most limiting factor of chloroplast CO_2_ delivery (including both stomatal conductance and mesophyll conductance components) and photosynthetic biochemistry ^151^, this phenomenon could be explained if stomata are either not limiting to plant photosynthesis under the study conditions or if secondary adjustments in leaf architecture compensate for changes in stomatal density ^150^. Future research efforts will be required to further elucidate the complex interplay between stomata, leaf architecture and plant physiology.

In addition to the lack of effect of reduced stomatal density on leaf photosynthesis, CO_2_ assimilation was also not impacted by the expression of SmNOX under the control of the CAB3 promoter. Thus, whilst *Sm*NOX was able to perturb NAD^+^ levels to mediate a reduction in stomatal density, it was not sufficient to impair photosynthesis. The reason for this is likely that the large pool sizes of NADH (24 – 55 μM) and NADPH (140 – 180 μM) ^131^ and the extraordinary high rates of photosynthetic NADPH production (∼70 μmol g^−1^ s^−1^) ^132^ would together mask the effect of *Sm*NOX activity in the mesophyll tissue. By contrast, epidermal cells contribute minimally toward total plant photosynthesis and contain small chloroplasts with high stroma to grana ratio which would produce much lower rates of NADPH ^152,153^. Thus, although *Sm*NOX is expressed to a high level throughout diverse photosynthesis tissues using the CAB3 promoter, the impact of this enzyme on cellular NAD^+^ status would be largest in stomatal progenitor cells.

Consistent with the lack of impairment of stomatal density or *Sm*NOX expression on photosynthetic CO_2_ assimilation, transgenic plants did not exhibit any difference in plant growth or biomass accumulation under well-watered conditions. However, increased vegetative growth and biomass accumulation was observed under water-deficit conditions. This improved tolerance to adverse water conditions is caused by the more conservative water-input requirements of these plants, driven by both the reduced rates of transpiration and improvement in photosynthetic water-use efficiency and is in agreement with other analyses of reduced stomatal density mutants ^71–73,75^. Moreover, the reduced leaf mass per area measured in *Sm*NOX-expressing lines would also favour biomass accumulation as this requires a lower resource investment per unit leaf tissue produced. Despite this improved tolerance to adverse water conditions, all *Sm*NOX-expressing lines exhibited a lower overall seed yield. Similar impairments in plant reproductive traits were also reported in NAD^+^ carrier mutants ^82,154^ and are likely caused by the role of NAD^+^ in seed ^155,156^ and pollen development ^157–160^. It would be interesting to investigate in future work whether this yield impairment is unavoidable, or whether it could be instead mitigated by use of other promoters with reduced off-target expression effects.

## Conclusion

As global food security is increasingly threatened by climate change, developing more climate-resilient crops presents an important goal for crop breeding and engineering efforts. Altering stomatal density represents a promising avenue to optimise the relationship between plant carbon assimilation, water demand, growth, and yield. Here, we provide a novel metabolic engineering strategy which can be exploited to reduce plant stomatal density by altering the endogenous NAD^+^ status of stomatal precursor cells. We demonstrate that this approach produces plants with improved photosynthesis-water relations and improved productivity under water-deficit conditions without any apparent cost on photosynthetic carbon assimilation or growth under normal conditions. Thus, this approach reveals a potential novel way in which plants can be altered to enhance climate resilience.

## Materials and Methods

### Gene accession numbers

The following list comprises the accession numbers of the genes used in this study: *Sm*NOX2 (WP_002268749.1); *At*RBCS1A (At1g67090); *At*PSBQA (At4g21280)*; At*OEP9 (At1g16000).

### Gene synthesis

A variant of the NAD(P)H oxidase (NOX) enzyme encoded by the Streptococcus mutans *nox2* gene but containing amino acid substitutions D192A/V193R/V194H/A199R was chosen for use in this study ^127^. This gene (hereinafter, termed *Sm*NOX) was codon optimised for expression in *Arabidopsis thaliana* and domesticated to remove the type IIS restriction enzymes BsaI or BpiI. Appropriate flanking regions compatible with Golden Gate Assembly ^161^ were added to the 5’ and 3’ regions, and the resulting coding sequence was commercially synthesized ^161^ (TWIST Bioscience, USA). The full-length coding sequence of *AtOEP9* and *At*RBCS1A, as well as a chloroplast stromal transit peptide from *AtPSBQA* (containing the first 87 amino acids of *AtPSBQA* including the intact stromal import domain but lacking a complete thylakoid targeting motif ^162^) were subject to these same steps for codon optimisation, domestication, addition of compatible flanking regions for Golden Gate assembly ^161^, and gene synthesis. The full-length sequences of each these genes are found in Supplemental File 2.

### Construct design and assembly

Genetic constructs for stable expression in plants were generated by cloning the coding sequence of *Sm*NOX downstream of the *A. thaliana* chlorophyll a/b-binding protein 3 (*At*CAB3; pTEI071) and upstream of the *A. tumefaciens* nopaline synthase terminator (nosT; piCH41421) in the Level 1 backbone position forward 2 vector pICH47742. Compositional forms of this Level 1 module were also generated in which *Sm*NOX was translationally fused at the N-terminus to either the *At*PSBQA transit peptide or the *At*OEP9 full-length coding sequence, respectively. Each of these constructs were subsequently assembled into Level 2 Golden Gate modules using the plant transformation vector acceptor backbone pAGM37443 ^163^ with the pFAST-R seed-coat specific fluorescent selection cassette ^164^ (pICSL70008) and the end Linker position 3 insert (pICH49266).

Genetic constructs for transient expression in protoplasts were generated by cloning the coding sequence of *Sm*NOX translationally fused to a C-terminal enhanced green fluorescent protein (GFP; pJOG176) downstream of the cauliflower mosaic virus 35S promoter (CaMV35S; EC15058) and upstream of the *A. tumefaciens* nosT (piCH41421) in the Level 1 backbone position forward 2 vector pICH47742. As above, compositional forms of this were also generated in which *Sm*NOX was translationally fused at the N-terminus to the *At*PSBQA transit peptide or the *At*OEP9 full-length coding sequence, respectively. Positive control constructs were also cloned using the same promoter, terminator, and backbone vector but containing only the coding sequence of the GFP (cytosolic fluoresce control) or the coding sequence of GFP translationally fused at the N-terminus to the full-length *At*RBCS1A gene sequence (chloroplast stroma fluoresce control). All cloning reactions were performed following the Golden Gate one-step one-pot protocol ^165^ as described in Supplemental File 1. The provenance of commercial vectors used during cloning are also provided in Supplemental File 1. Full-length sequences of each expression cassette generated is provided in Supplemental File 2. The full set of oligonucleotides used for cloning, colony PCR, and Sanger sequencing are found in Supplemental File 3. The thermocycler conditions used for all PCR reactions are also described in Supplemental File 3.

### Protoplast isolation, transformation, and visualisation

Mesophyll protoplasts were isolated from leaves of 4-week old wild-type *Arabidopsis thaliana* Col-0 plants following the ‘Tape-*Arabidopsis* Sandwich’ method ^167^. Protoplast transformation was performed using the calcium/polyethylene glycol-mediated method ^168^, as modified in ^167^. Transformed protoplasts were incubated in the dark overnight at 25°C prior to imaging. Cell fluorescence was visualised using a Zeiss LSM 880 Airy Scan confocal microscope. Zeiss ZEN software (Zeiss) was used for confocal laser scanning microscopy and subsequent image processing.

### Plant material and stable transformation

Stable transformation of *A. thaliana* ecotype Columbia (Col-0) background was performed using the agrobacterium-mediated floral dip method ^169^. Positive transformants were initially screened based upon seed fluorescence visualised by epifluorescence microscopy (Leica M165 stereo microscope), and confirmed by genotyping PCR and gel electrophoresis using the OneTaq® 2X Master Mix with Standard Buffer (NEB, Catalog #M0482S) following manufacturer’s instructions for reaction set-up and the oligonucleotides and thermocycler conditions described in Supplemental File 3. Single insertion events were subsequently identified by TaqMan® real-time PCR with fluorescent probes and primers which bind to the TagRFP selectable marker (Idna GENETICS) and homozygous individuals were isolated. For the above purposes of genotyping PCR and TaqMan® real-time PCR, genomic DNA was isolated from mature leaves of T1 plants using the cetyl trimethylammonium bromide (CTAB) method ^170^. In total, three single-copy homozygous lines were taken forward as the basis of the analysis herein for each *Sm*NOX-expressing transgenic plant (cyto-*Sm*NOX, stromal-*Sm*NOX and CIMS-*Sm*NOX). All experiments described are performed on T3 and T4 generation plants.

### Stomatal density measurements

Epidermal impressions from the abaxial surface of healthy and fully-expanded excised leaves of 8-week old plants were obtained following the clear nail polish method ^171^. All leaves used for this analysis were of similar size. After mounting on microscope slides, stomatal density was measured using a Leica DMR8 light microscope at ×20 magnification under brightfield view. For each sample, stomatal density counts were calculated across five independent fields of view of the same leaf and across 9 – 10 biological replicates per line.

### Leaf mass area measurements

A leaf puncher was used to harvest a fixed area of tissue from healthy and fully-expanded leaves of 8-week-old plants. Leaf dry mass was subsequently determined after 48 hours incubation at 65 °C. All leaves were of similar size and were harvested in the middle of the photoperiod. Leaf mass area (LMA, gLJm^−2^) was subsequently determined as the fixed leaf area divided by leaf dry mass.

### In vitro enzyme assays

A consistent area of leaf tissue was harvested from healthy and fully-expanded leaves of 6-week-old plants using a leaf puncher during the middle of the photoperiod. Leaf material was flash frozen in liquid nitrogen, ground using metal beads and a tissue lyser (Thermo Scientific, TissueLyser II), and resuspended in 500 μL ice cold extraction buffer (50 mM potassium phosphate, 5 mM MgCl2, 1 mM EDTA, 3% Triton X-100) with 1x protease inhibitor cocktail (Sigma, Catalog #P9599-5ML). Homogenate was centrifuged for 1 minute at 12,000 rpm at 4°C, and total crude protein extract was isolated from supernatant.

Enzyme assays were performed following the method described in ^127^ with some alterations. In brief, all reactions were performed using a total volume of 200 μL in unsealed 96 well plate at 25°C. Typical reaction mixtures contained 50 mM potassium phosphate buffer (pH 7.5) and 250 μM of either NADH or NADPH, were vortexed for saturation with oxygen in air (250 μM), and were initiated by addition of 30 μL undiluted crude leaf lysate. Negative controls containing ddH_2_O in the place of leaf lysate were included in every assay. Consumption of NADH or NADPH was monitored spectrophotometrically at 340 nm using a plate reader (BMG Labtech FLUOstar Omega). Each biological replicate was repeated over two technical replicates wells and rates of enzyme reaction were calculated over the linear part of the curve over a duration of at least 5 minutes. Reaction rates were normalized per unit NADH/NADPH oxidation by standard curve.

### Leaf gas exchange

Gas exchange analyses were performed using a LI-6800 open-path gas exchange system equipped with a 6800-01A fluorometer head (LI-COR, USA) using the 2 cm^2^ leaf chamber and 90% red and 10% blue actinic light under normal 21% O_2_ air. The youngest fully-expanded leaf was used for each plant and all measurements were taken prior to onset of flowering. Downstream statistical analysis, model fitting, and parameter estimation from leaf gas exchange data was performed as described in Supplemental File 1.

Response of plant gas exchange to light followed the method of ^172^ with some alterations. In brief, plants were first adapted at 2,200 μmol m^−2^ s^−1^ for 20 minutes, and photosynthesis was then measured at step-wise intervals of 2200, 2000, 1650, 1300, 1000, 750, 500, 400, 300, 200, 150, 100, 50, 0 and 2200 μmol m^−2^ s^−1^ light. The environment was held constant at 400 μmol mol^−1^ CO_2_, 30°C leaf temperature, 55% relative air humidity, 500 µmol s^−1^ flow rate, and 10,000 rpm fan speed. No matching was performed during the analysis and a 4-minute wait time was used between each light condition.

For the response of plant gas exchange to intercellular CO_2_, plants were first adapted at 400 μmol mol^−1^ CO_2_ for 20 minutes and photosynthesis was then measured at step-wise intervals of 400, 0, 50, 75, 100, 200, 400, 800, 1200, 1600, 400 μmol mol^−1^ reference atmospheric CO_2_ concentrations. The environment was held constant at 1500 µmol m^−2^ s^−1^ light, 30°C leaf temperature, 55% relative air humidity, 500 µmol s^−1^ flow rate, and 10,000 rpm fan speed. Matching of CO_2_ and H_2_O was performed before each measurement with a minimum and maximum wait time of 3 and 4 minutes, respectively.

Investigation of stomatal kinetics in response to step changes in light followed the method of ^142^. In brief, plants were first adapted to 100 μmol m^−2^ s^−1^ light for 60 minutes and were then subject to a step increase to 1000 μmol m^−2^ s^−1^ for 60 minutes followed by a step decrease back to 100 μmol m^−2^ s^−1^ for 30 minutes. The environment was held constant at 400 μmol mol^−1^ CO_2_, 30°C leaf temperature, 1.8 kPa vapour pressure deficit, 500 µmol s^−1^ flow rate, and 10,000 rpm fan speed. No matching was performed during the analysis and measurements were taken every minute.

For the response of plant gas exchange to temperature, plants were first adapted at 20°C for 20 minutes and photosynthesis was then measured at step-wise intervals of 20, 25, 30, 35, 40 and 45°C reference air temperatures. The environment was held constant at 1500 µmol m^−2^ s^−1^ light, 400 μmol mol^−1^ CO_2_, 500 µmol s^−1^ flow rate, and 10,000 rpm fan speed. As previously described in the thermal analysis of ^173^, it is difficult to maintain VPD at temperatures above 30°C. Thus, leaf vapour pressure deficit (VPD) was maintained at 1.5 kPa for measurements at 20, 25 and 30°C, 2.5 kPa for the measurement at 35°C, 3.5 kPa for the measurement at 40°C and 4.5 kPa for the measurement at 45 °C. Matching of CO_2_ and H_2_O was performed before each measurement with a minimum and maximum wait time of 10 and 14 minutes, respectively.

### Plant growth conditions

Seeds were grown in individual pots directly on soil (Levington Seed modular compost) and were stratified for 2 days at 4°C to overcome dormancy. Unless explicitly stated, plants were grown in climate-controlled environment conditions in floor-standing growth chambers (Aralab FitoclimaD1200 PLH) under short-day conditions (8-hour light/16-hour dark cycle) at 21°C, ambient 400 ppm CO_2_, 50% relative humidity, and 125 μmol m^−2^ s^−1^ light and were maintained under well-watered conditions.

### Plant growth assays

For analysis of plant growth, after stratification plants were transferred to controlled environment chambers (CERs) at The University of Oxford and grown under a 16-hour/8-hour day/night period, 21°C, and 250 μmol m^−2^ s^−1^ light. To minimize confounding and positional effects, plant genotypes were grown in a fully randomised block design and the position of trays were rotated every 2 days. For well-watered experiments, plants were watered on a weekly basis as usual from the base of the tray. For water-deficit growth experiment, plant trays were watered prior to stratification as above, but water was then withheld completely for 15 days after germination to maintain sub-optimal and deteriorating soil-water saturation levels during vegetative growth. After this period, watering was resumed following the method described in the well-watered growth experiments.

Aerial photographs were taken daily during vegetative growth. Image processing was performed to correct for orientation and perspective using Yet Another Scanning Wizard (YASW) (https://github.com/ImageProcessing-ElectronicPublications/yasw). Computational analysis was performed to derive growth parameters. Green pixels were isolated from images using an in-built script, and visible rosette area was computed with ImageJ software^174^ using the in-built ‘Analyze Particles’ function. Time of bolting was measured as the number of days after stratification when the inflorescence was greater than 1 cm in height above the rosette. Time of flowering time was measured as the number of days after stratification at which the first flower was open and petals were visible. Plant height was measured as the maximum height of the primary inflorescence from the base of the rosette at 32 days post stratification. Total seed weight was measured after allowing plants to completely dry for a month.

## Funding

This work was funded by the Royal Society and the European Union’s Horizon 2020 research and innovation program under grant agreement number 637765. JWB was funded by the BBSRC through BB/J014427/1. This research was funded in whole, or in part, by the BBSRC number BB/J014427/1. For the purpose of open access, the author has applied a CC BY public copyright license to any Author Accepted Manuscript version arising from this submission.

## Data Availability

All data used in this study is provided in the supplemental material.

## Author Contributions

JWB and SK conceived the study, designed the experiments, interpreted and analyzed the data, and wrote the manuscript. JWB conducted the experiments.

## Conflicts of interest

SK is co-founder of Wild Bioscience Ltd. SK and JWB have applied for a patent on the work presented in this study.

