## Supplemental File 1 for "Metabolic engineering of stomatal precursor cells enhances photosynthetic water-use efficiency and vegetative growth under water-deficit conditions in *Arabidopsis thaliana*"

### Supplemental Results

#### NOX expression results in tighter coordination between rates of assimilation and stomatal conductance

To further investigate the *iWUE* phenotype, the relative coordination between *A* and *g*_s_ was evaluated across the different light intensities. For this purpose, values of *g*_s_ were compared between those observed at *g*_s sat_ and the modelled light intensity where 95% maximum assimilation (*g_s_ _A95_*) was achieved (beyond which neither CO_2_ delivery nor light is considered limiting to photosynthesis). This revealed that rates of *g*_s_ in wild-type plants continue to increase by 8.1% after *g_s_ _A95_* is achieved without resulting in any benefit in terms of CO_2_ assimilation (Figure 3G, Supplemental File 1, Figure S6A – S6C, table S3). In contrast, the magnitude of this “overshooting” behaviour was significantly dampened in transgenic plants, where an increase in *g_s_* of only 3.3 – 4.4% was observed after *g_s_ _A95_* was attained (Figure 3G, Supplemental File 1, Figure S6A – S6C, table S3). This same result was recapitulated when comparing the respective light intensities at which 95% maximum assimilation rate (PPFD _A95_) and *g*_s sat_ (PPFD *g*_s sat_) were achieved. For instance, PPFD _A95_ and PPFD *g*_s sat_ were more similar to one another in transgenic (cyto-*Sm*NOX: 828.6 ± 161.3 μmol m^-2^ s^-1^, 156.2%; stromal-*Sm*NOX: 621.4 ± 142.8 μmol m^-2^ s^-1^, 124.6%; CIMS-*Sm*NOX: 847.6 ± 169.7 μmol m^-2^ s^-1^, 168.2%) compared to wild-type plants (1385.7 ± 235.5 μmol m^-2^ s^-1^, 290.0%) (Figure 3H, Supplemental File 1, Figure S6D – S6F, table S3). Thus, the reduced stomatal density caused by *Sm*NOX expression resulted in an enhanced water-use efficiency and tighter coupling of *g*_s_ and *A* across different light conditions.

### Supplemental Materials and Methods

#### Cloning and construct assembly

Cloning was performed using the Golden Gate one-step one-pot protocol ^1^ with modified thermocycler conditions consisting of 80 cycles of 5 minutes digestion (37°C) and 5 minutes ligation (16°C). All reactions contained T4 DNA ligase (NEB, Catalog #M0202S) and the Type IIS restriction enzymes BsaI-HF®v2 (Level 1 assembly; NEB, Catalog #R3733S) or BpiI (Level 2 assembly; Thermo Fisher Scientific, Catalog # ER1012). Assembled plasmids were checked by colony PCR and gel electrophoresis using the OneTaq® 2X Master Mix with Standard Buffer (NEB, Catalog #M0482S) using generic primers that bind to the vector backbone, and sequences of inserts were confirmed by Sanger sequencing (Source BioScience, UK). Full plasmid sequencing (SNPsaurus, USA) was performed on all Level 2 constructs prior to transformation. Where 5’ and 3’ overhangs of vectors were not compatible for a given reaction, gene inserts were amplified by PCR with altered overhangs and resulting linear fragments were used directly for cloning. All PCR reactions for cloning was performed using Q5® High-Fidelity DNA Polymerase (NEB, Catalog #M0491S) following manufacturer’s instructions for reaction set-up.

#### Vectors

pTEI071 and pJOG176 were a gift from Johannes Stuttmann (Addgene plasmid #105376; http://n2t.net/addgene:105376; RRID:Addgene_105376, and Addgene plasmid #105392; http://n2t.net/addgene:105392; RRID:Addgene_105392, respectively) ^2^**.** piCH41421 was a gift from Sylvestre Marillonnet and Nicola Patron (Addgene plasmid #50339; http://n2t.net/addgene:50339; RRID:Addgene_50339) ^3^. pICSL70008 was a gift from Nicola Patron (Addgene plasmid #50336; http://n2t.net/addgene:50336; RRID:Addgene_50336) ^3^. pICH47742 and pICH49266 were a gift from Sylvestre Marillonnet (Addgene plasmid # 48001; http://n2t.net/addgene:48001; RRID:Addgene_48001, and Addgene plasmid #48024 ;http://n2t.net/addgene:48024; RRID:Addgene_48024) ^4^**.** pAGM37443 was a gift from Sylvestre Marillonnet (Addgene plasmid #153218; http://n2t.net/addgene:153218; RRID:Addgene_153218) ^5^. EC15058 was a gift from Ben Miller.

#### Statistical analysis, model fitting, and parameter estimation of leaf gas exchange data

For statistical analysis of the response of plant gas exchange to light, non-linear curves were fit to the data by implementing the non-rectangular hyperbola model ^6^ using the ‘fit_photosynthesis’ function in the R package photosynthesis V2.1.4 ^7,8^. Values of light compensation point, mitochondrial respiration in the light, apparent quantum yield, and light-saturated electron transport rate were derived from these fitted models using this function. Light-saturated rates of assimilation (*A*_sat_), stomatal conductance, and intrinsic water-use efficiency were manually determined as the highest value of these respective traits across the light response curve. The 95% maximum assimilation rate was also manually from these models.

For statistical analysis of the response of plant gas exchange to CO_2_, the Farquhar- von Caemmerer- Berry model of leaf photosynthesis was fit to the data using the ‘fitaci’ function from the R package plantecophys ^9^. For this purpose, the bilinear method was used with triose phosphate utilization implemented to improve model fitting. Values of the maximum carboxylation rate of rubisco (*V*_cmax_), maximum electron transport rate (*J*_max_), CO_2_ compensation point, the Michaelis constant of rubisco for CO_2_ in 21% O_2_ air, and the CO_2_ concentration at the transition between rubisco-limited and ribulose-1,5-bisphosphate regeneration-limited photosynthesis were all derived from these fitted models using this function. Estimates of *V*_cmax_ and *J*_max_ re-scaled to 25°C were also generated from in-built temperature corrections of the data using the ‘fitaci’ function ^9^. The CO_2_–saturated assimilation rate (*A*_max_) was manually determined as the highest value of this trait across the CO_2_ response curve. The CO_2_ concentration at the transition between RuBP regeneration-limited photosynthesis and triose-phosphate utilization-limited photosynthesis were not reported as could not be estimated in all cases.

Statistical analysis of stomatal kinetic parameters to step changes in light intensity between 100 μmol m^−2^ s^−1^ to 1000 μmol m^−2^ s^−1^ were derived using a dynamic sigmoidal mathematical model ^10,11^. For this purpose, model fitting was implemented using the licornetics V2.1.2 source code downloaded from the github repository <https://github.com/lbmountain/licornetics>. Estimates of the maximal rate of stomatal opening, time taken to achieve new steady-state stomatal conductance, and the initial lag in response time of *g*_s_ to the step increase in light intensity from 100 μmol m^−2^ s^−1^ to 1000 μmol m^−2^ s^−1^, as well as the estimated time taken to achieve new steady-state stomatal conductance after the step decrease in light intensity from 1000 μmol m^−2^ s^−1^ to 100 μmol m^−2^ s^−1^ were all derived from models fitted in this way.

### Supplemental Figures

#### Supplemental File 1, Figure S1

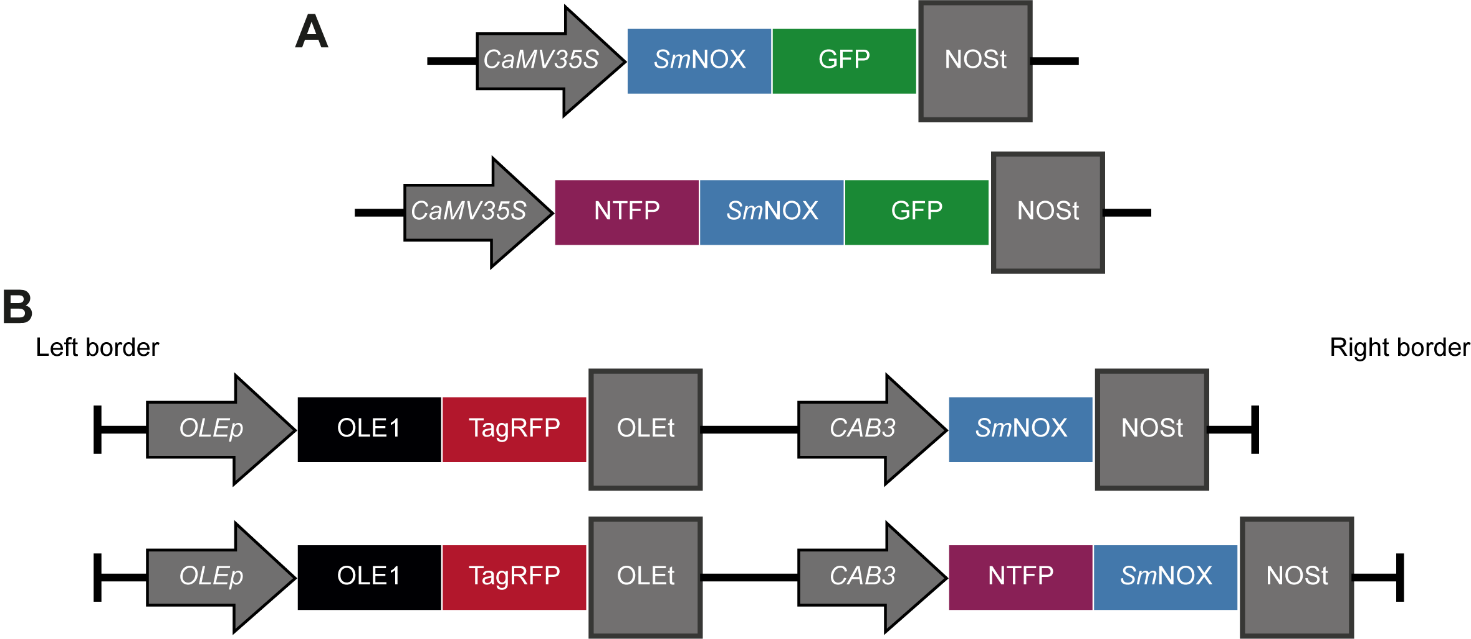

**Figure S1.** *Sm*NOX expression cassettes used for experiments in *Arabidopsis thaliana*. A) Simplified schematic of the expression cassette used for *Sm*NOX transient expression in *A. thaliana* protoplasts. The construct contains a CaMV 35S promoter (CaMV35S) and a nopaline synthase terminator (NOSt) controlling the expression of the *Sm*NOX coding sequence translationally fused to eGFP at the C-terminus. Top: untargeted *Sm*NOX. Bottom: *Sm*NOX targeted by an N-terminal fusion peptide (NTFP). NTFPs are derived from *A. thaliana* genes and include the predicted transit peptide from the photosystem II subunit QA gene or the full-length outer envelope protein 9 gene. B) As in (A), but a simplified schematic of the expression cassette used for *Sm*NOX stable expression in *A. thaliana*. In contrast to (A), *Sm*NOX does not contain a C-terminal eGFP and is driven under the control of the chlorophyll a/b-binding protein 3 (CAB3) promoter. All constructs also include an additional expression cassette encoding a seed coat specific fluorescent selectable marker which consists of monomeric TagRFP from *Entacmaea quadricolor* fused to the coding sequence of the *Arabidopsis thaliana* oleosin1 gene (OLE1) under the control of the native oleosin1 promoter (OLEp) and terminator (OLEt), respectively. The complete nucleotide and amino acid sequences for each expression cassette can be found in Supplemental File 2

#### Supplemental File 1, Figure S2

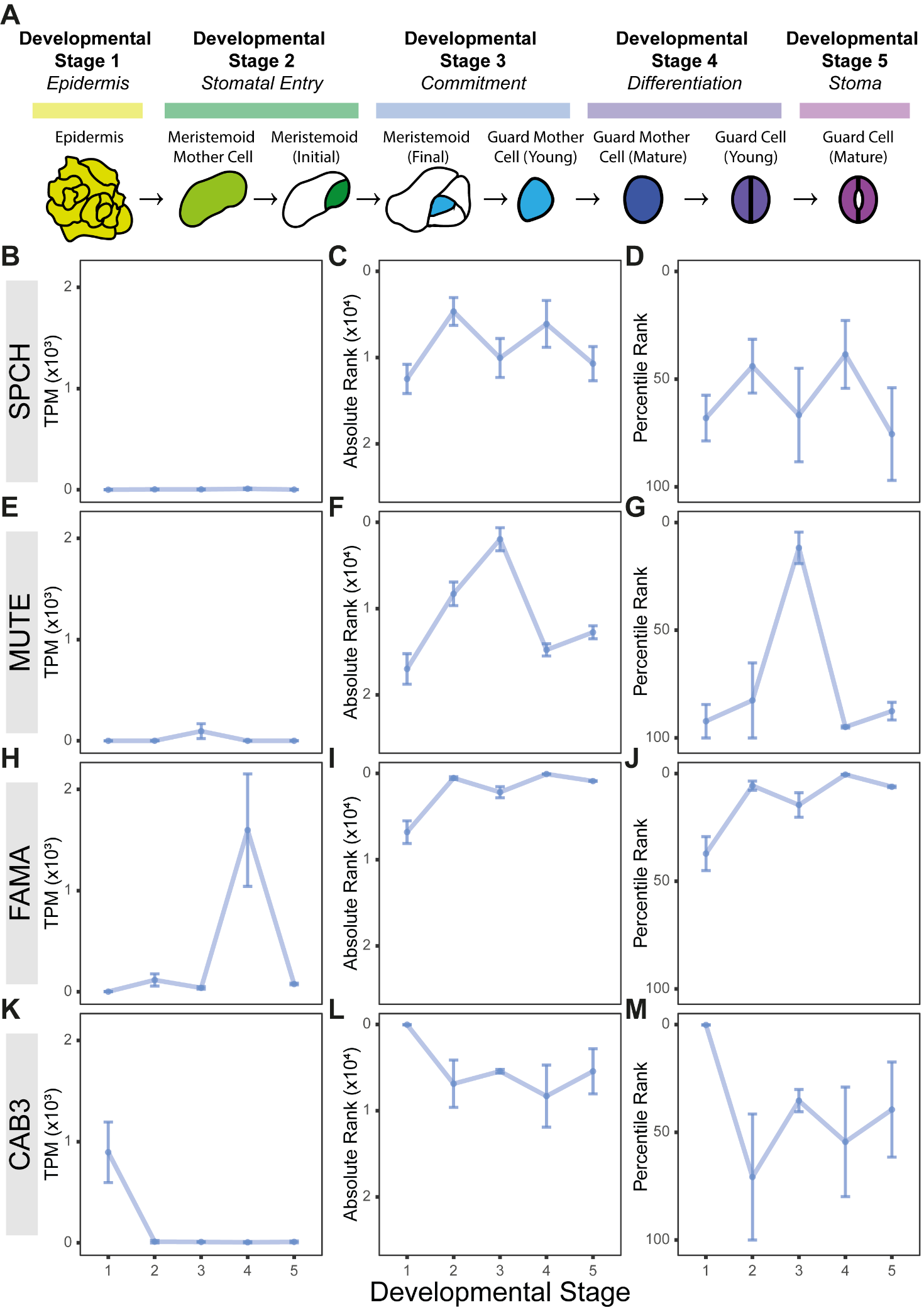

**Figure S2.** The chlorophyll a/b binding protein (CAB3) promoter drives high gene expression in stomatal precursor cells. A) Schematic of the developmental trajectory of stomatal lineage differentiation including all major steps and fate transitions from undifferentiated epidermal cell (Developmental Stage 1), to stomatal lineage entry (Developmental Stage 2), stomatal commitment (Developmental Stage 3), stomatal differentiation (Developmental Stage 4), and finally, development of a fully-functional stoma (Developmental Stage 5). Figure adapted from ^12^. B) Temporal profile of SPEECHLESS (SPCH) transcript abundance (TPM; Transcripts Per Million) along the stomatal developmental trajectory in leaves of *Arabidopsis thaliana*. Respective developmental stages follow those described in (A). Representative transcriptomes of populations of cells at each developmental stage were obtained from ^12^. C) As in (B), but for the transcript abundance of SPCH expressed as a rank against the distribution of the transcript abundances of all other nuclear-encoded genes in *A. thaliana*. Ranks are calculated in descending order of gene transcript levels (i.e., low ranks denote genes with the highest relative expression and *vice versa*) and the y-axis has been inverted for ease of visualisation. D) As in (B), but for the transcript abundance of SPCH expressed as a percentile rank against the distribution of the transcript abundances of all other nuclear-encoded genes in *A. thaliana*. Gene percentile ranks are calculated in descending order of gene transcript levels (i.e., low percentile ranks denote genes with the highest relative expression and *vice versa*) and the y-axis has been inverted for ease of visualisation. E – G) As in (B), (C), and (D), but for MUTE. H – J) As in (B), (C), and (D), but for FAMA. K – M) As in (B), (C), and (D), but for CAB3. The raw data can be found in Supplemental File 5.

#### Supplemental File 1, Figure S3

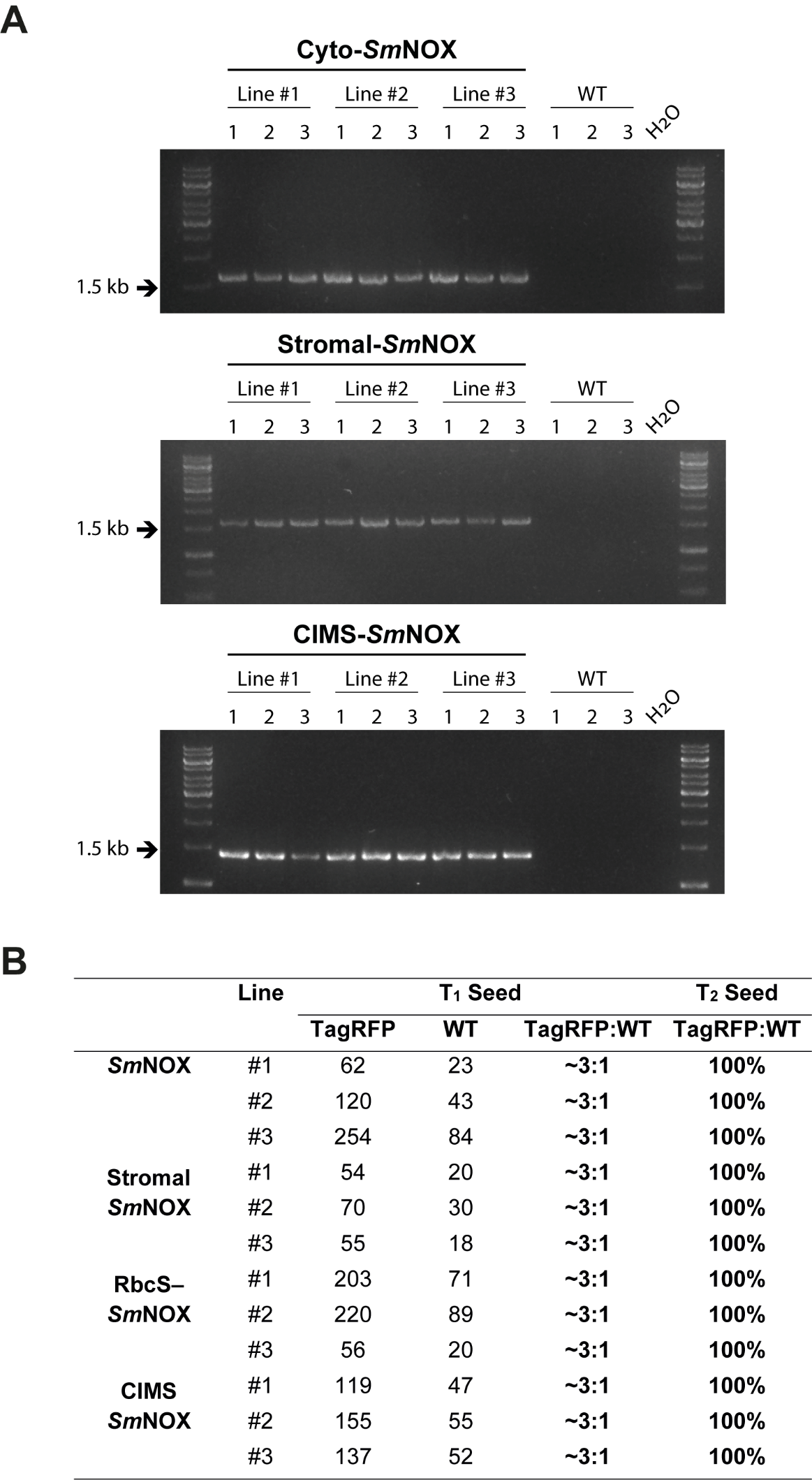

**Figure S3.** Verification of *Sm*NOX implementation. A) Genotyping of transgenic T_3_ *Sm*NOX and wild-type (WT) plants based on primers specific for each gene insertion. The primers, reaction set up, and thermocycler conditions used for polymerase chain reaction (PCR) can be found in Supplemental File 3. B) Segregation analysis data of seed from T_1_ and T_2_ generation plants.

#### Supplemental File 1, Figure S4

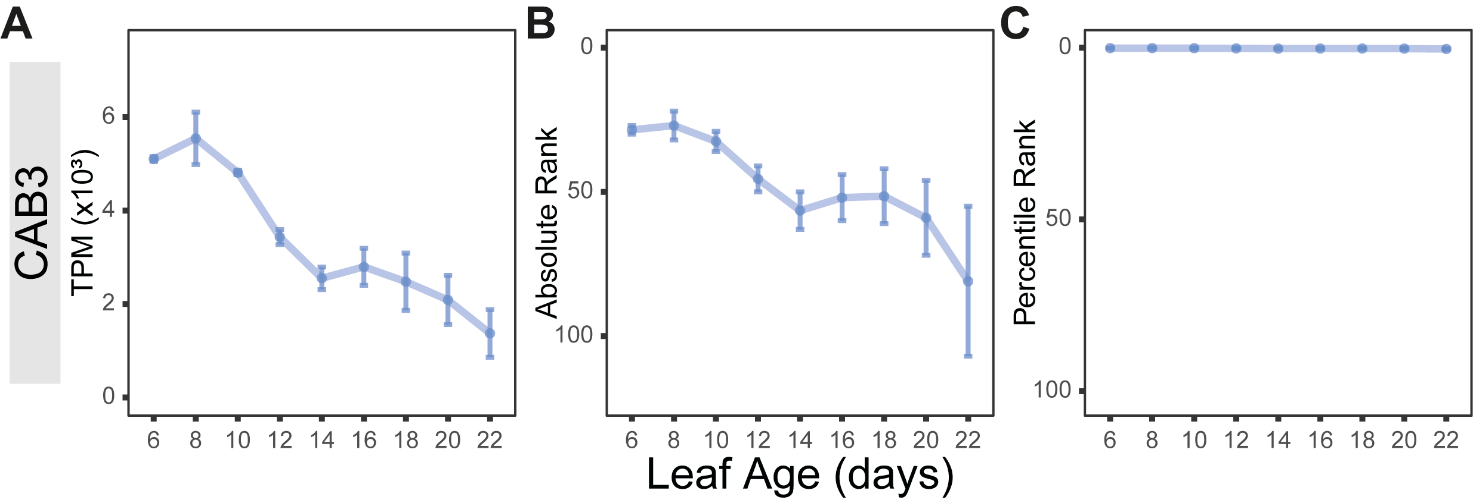

**Figure S4.** The chlorophyll a/b binding protein (CAB3) promoter drives highest levels of gene expression during early leaf development. A) Temporal profile of CAB3 transcript abundance (TPM; Transcripts Per Million) along a gradient of leaf aging in *Arabidopsis thaliana*. Representative transcriptomes from leaves at each age were obtained from ^13^. C) As in (B), but for the transcript abundance of the CAB3 gene expressed as a rank against the distribution of transcript abundances of all other nuclear-encoded genes in *A. thaliana*. Gene ranks are calculated in descending order of gene transcript levels (i.e., low ranks denote genes with the highest relative expression and *vice versa*) and the y-axis has been inverted for ease of visualisation. D) As in (B), but for the transcript abundance of the CAB3 gene expressed as a percentile rank against the distribution of all other nuclear-encoded genes in *A. thaliana*. Gene percentile ranks are calculated in descending order of gene transcript levels (i.e., low percentile ranks denote genes with the highest relative expression and *vice versa*) and the y-axis has been inverted for ease of visualisation. The raw data can be found in Supplemental File 5.

#### Supplemental File 1, Figure S5

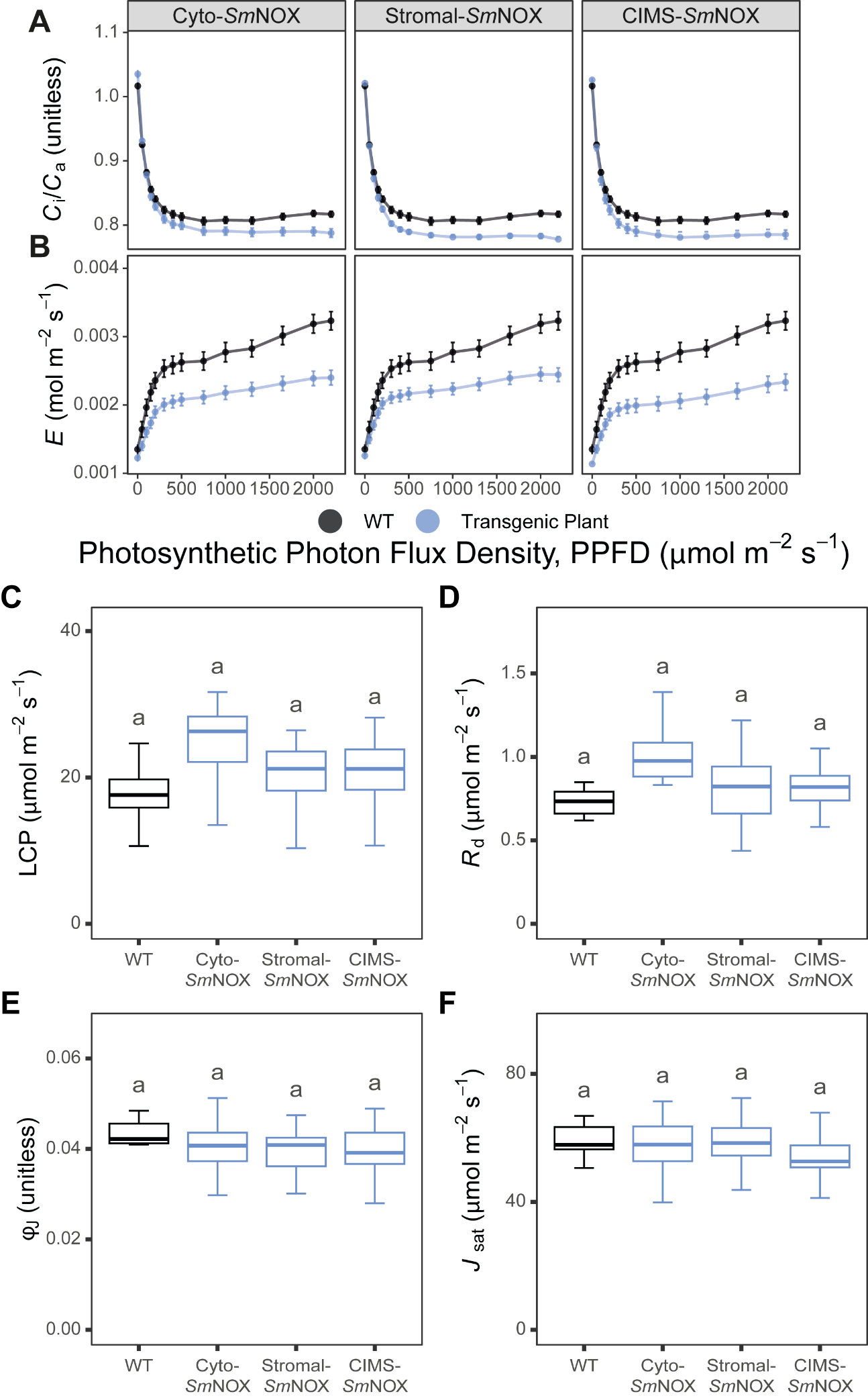

**Figure S5.** Additional parameters in the response of plant gas exchange to light (photosynthetic photon flux density: PPFD, μmol m^-2^ s^-1^). A) The ratio of intercellular CO_2_ to ambient CO_2_ (*C*_i_/*C*_a_, unitless) at different PPFD in wild-type (black) and transgenic plants (blue). WT: wild-type. Cyto-*Sm*NOX: cytosolic *Sm*NOX. Stromal-*Sm*NOX: chloroplast stromal *Sm*NOX. CIMS-*Sm*NOX: chloroplast intermembrane space *Sm*NOX. Data represent mean ± 1 S.E. B) as in (A), but for plant transpiration rate (*E*, mol m^−2^ s^−1^). C) Boxplot depicting the light compensation point (LCP, μmol mol^−1^). The colour scheme and abbreviations follow that described in (A). D) As in (C), but for the mitochondrial respiration in the light (*R*_d_, μmol m^−2^ s^−1^). E) As in (C), but for the modelled apparent quantum yield (ΦJ, μmol m^−2^ s^−1^). F) As in (C), but for the light-saturated electron transport rate (*J*sat, μmol m^−2^ s^−1^). All data represent an average across three independent single copy lines (*n* = 7 – 8 per individual line). Differences between transgenic plants and WT are assessed by Fisher LSD post-hoc analysis following a two-way ANOVA, where letters above each box represent statistically significant differences in mean values (p ≤ 0.05). The raw data can be found in Supplemental File 4.

#### Supplemental File 1, Figure S6

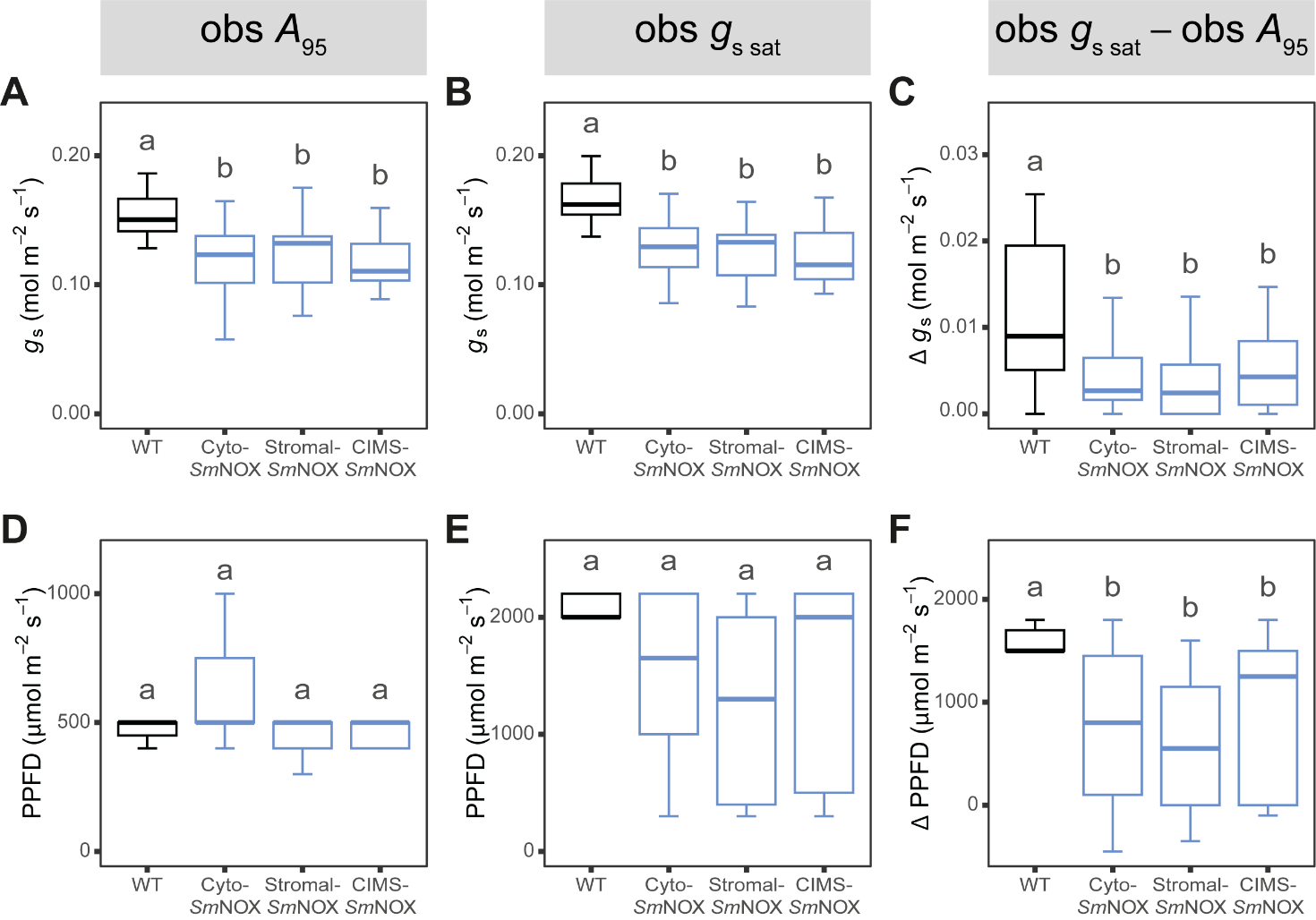

**Figure S6.** The relative coordination between rates of plant photosynthesis and gas exchange across light response curves. A) Boxplot depicting the stomatal conductance at the light intensity at which 95% maximum assimilation rate is achieved (*g_s_ _A95_*, mol m^−2^ s^−1^) in wild-type (black) and transgenic plants (blue). WT: wild-type. Cyto-*Sm*NOX: cytosolic *Sm*NOX. Stromal-*Sm*NOX: chloroplast stromal *Sm*NOX. CIMS-*Sm*NOX: chloroplast intermembrane space *Sm*NOX. B) As in (A), but for the light-saturated stomatal conductance (*g*_s sat_, mol m^−2^ s^−1^). C) As in (A), but for difference in stomatal conductance between the light intensity at which 95% maximum assimilation rate and the light-saturated stomatal conductance is achieved (Δ *g*_s_, mol m^-2^ s^-1^). D) As in (A), but for the light intensity at which 95% maximum assimilation rate is achieved (PPFD _A95_, μmol m^−2^ s^−1^). E) As in (A), but for the light intensity at which light-saturated stomatal conductance is achieved (PPFD *_g_*_s sat_, μmol m^−2^ s^−1^). F) As in (A), but for the difference in irradiance between the light intensity at which 95% maximum assimilation rate and the light-saturated stomatal conductance is achieved (Δ PPFD, μmol m^-2^ s^-1^). All data represent an average across three independent single copy lines (*n* = 7 – 8 per individual line). Differences between transgenic plants and WT are assessed by Fisher LSD post-hoc analysis following a two-way ANOVA, where letters above each box represent statistically significant differences in mean values (p ≤ 0.05). The raw data can be found in Supplemental File 4.

#### Supplemental File 1, Figure S7

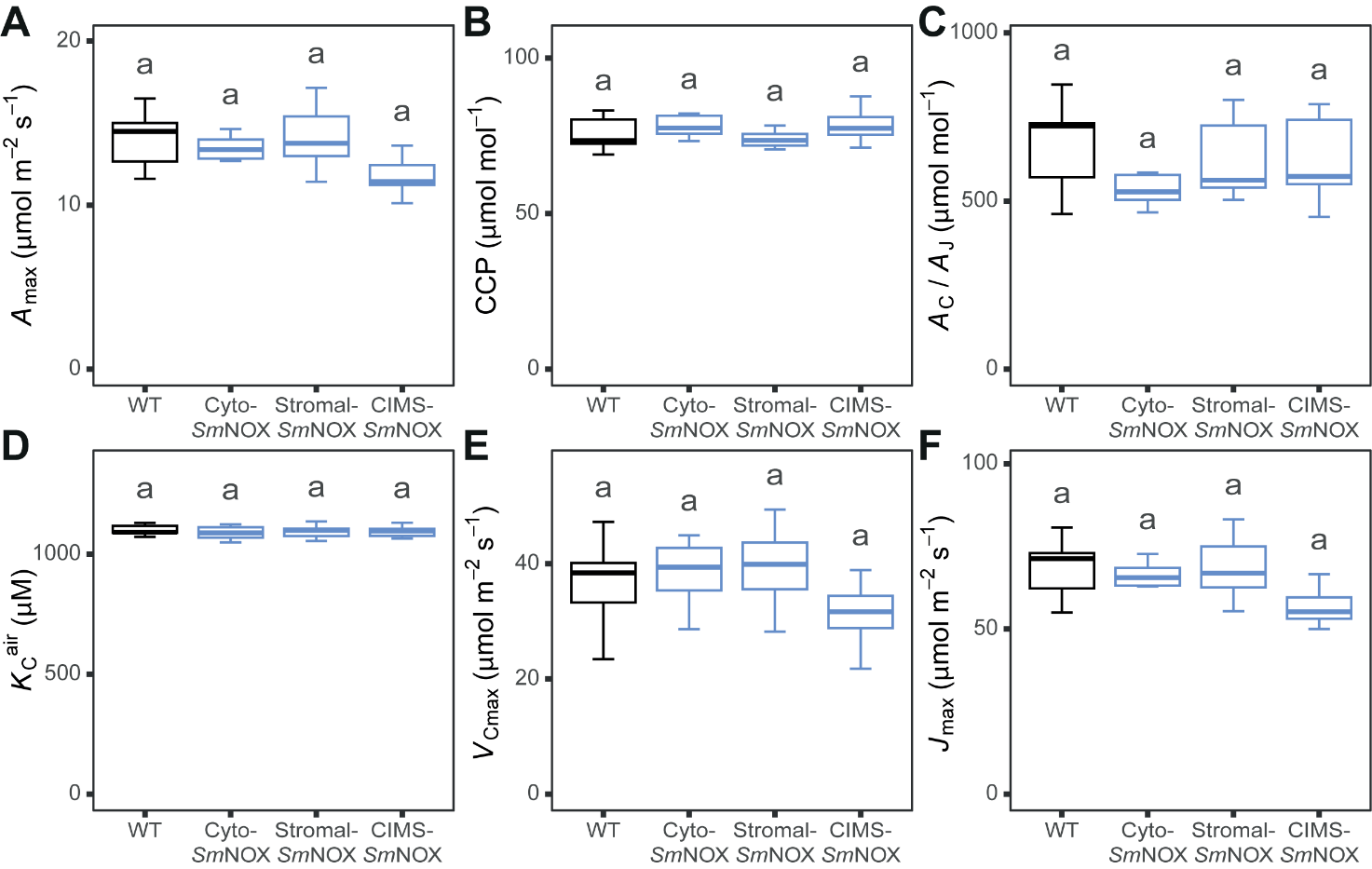

**Figure S7.** The response of plant gas exchange to intercellular CO_2_ concentration in wild-type (black) and transgenic plants (blue). WT: wild-type. Cyto-*Sm*NOX: cytosolic *Sm*NOX. Stromal-*Sm*NOX: chloroplast stromal *Sm*NOX. CIMS-*Sm*NOX: chloroplast intermembrane space *Sm*NOX. A) Boxplot depicting the CO_2_–saturated assimilation rate (*A*_max_, μmol m^−2^ s^−1^). B) As in (A), but for the estimated CO_2_ compensation point (CCP, µmol mol^-1^). C) As in (A), but for the estimated CO_2_ concentration at the transition between rubisco-limited and ribulose-1,5-bisphosphate (RuBP) regeneration-limited photosynthesis (*A*_C_/*A*_J_, µmol mol^-1^). D) As in (A), but for the estimated Michaelis constant of rubisco for CO_2_ in 21% O_2_ air (*K*_C_^air^, µM). E) As in (A), but for the estimated maximum carboxylation rate of rubisco at 30°C (*V*_cmax_, µmol m^−2^ s^−1^). F) As in (A), but for the estimated maximum electron transport rate at 30°C (*J*_max_, µmol e^−^ m^−2^ s^−1^). All data represent an average across three independent single copy lines (*n* = 3 – 5 plants per line). Differences between transgenic plants and WT are assessed by Fisher LSD post-hoc analysis following a two-way ANOVA, where letters above each box represent statistically significant differences in mean values (p ≤ 0.05). The raw data can be found in Supplemental File 4.

#### Supplemental File 1, Figure S8

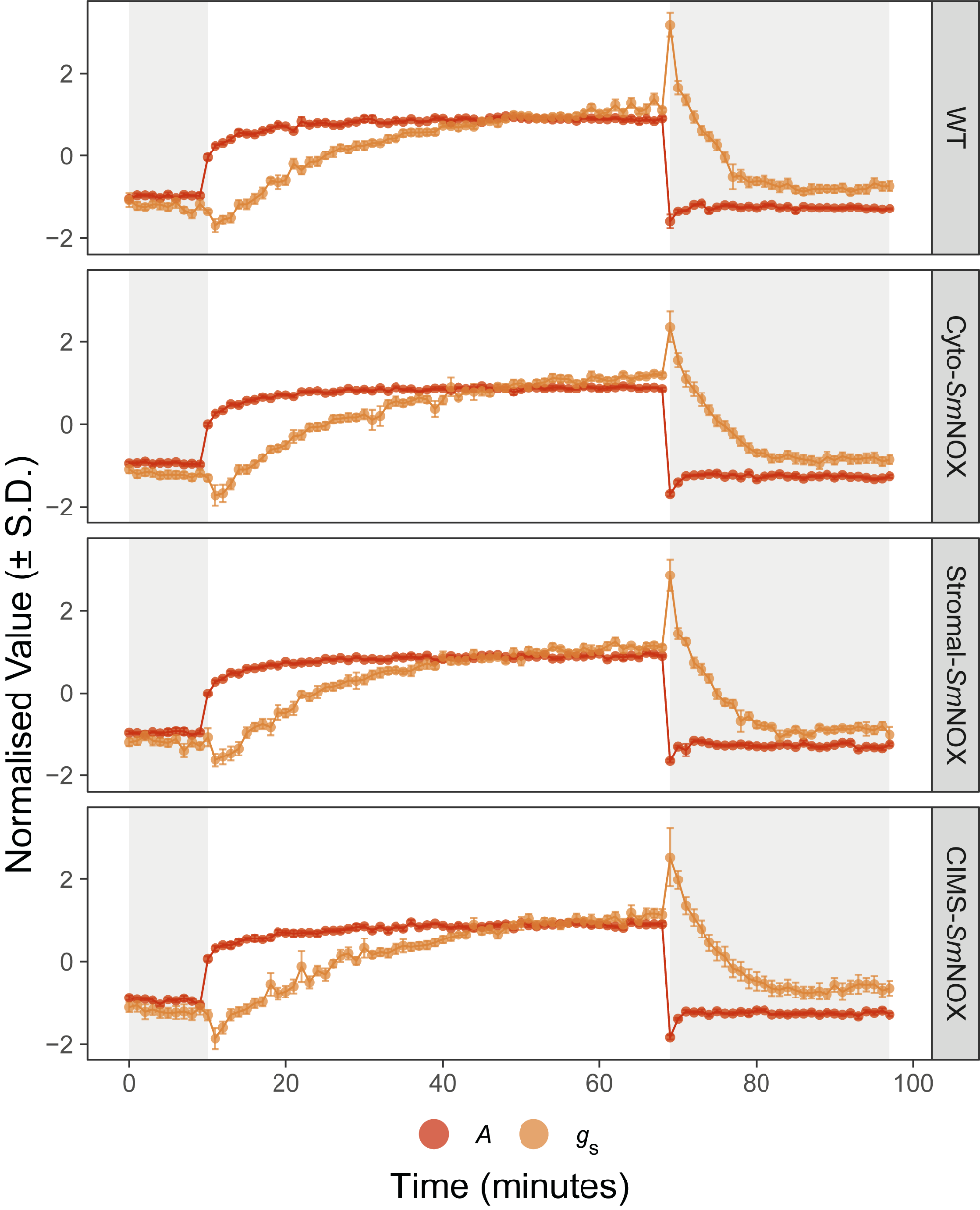

**Figure S8.** The normalised temporal response of plant CO_2­_ assimilation rate (*A*, μmol m^−2^ s^−1^; red) and stomatal conductance (*g*_s,_ mol m^−2^ s^−1^; orange) to step changes in light intensity. Shaded and unshaded areas represent periods of light intensity of 100 μmol m^−2^ s^−1^ and 1000 μmol m^−2^ s^−1^, respectively. WT: wild-type. Cyto-*Sm*NOX: cytosolic *Sm*NOX. Stromal-*Sm*NOX: chloroplast stromal *Sm*NOX. CIMS-*Sm*NOX: chloroplast intermembrane space *Sm*NOX. For each parameter, data are normalised by the standard deviation away from the mean calculated across the entire temporal response of that individual. Data represent mean ± 1 S.E of normalised values. All data represent an average across three independent single copy lines (*n* = 3 – 4 per individual line). The raw data can be found in Supplemental File 4.

#### Supplemental File 1, Figure S9

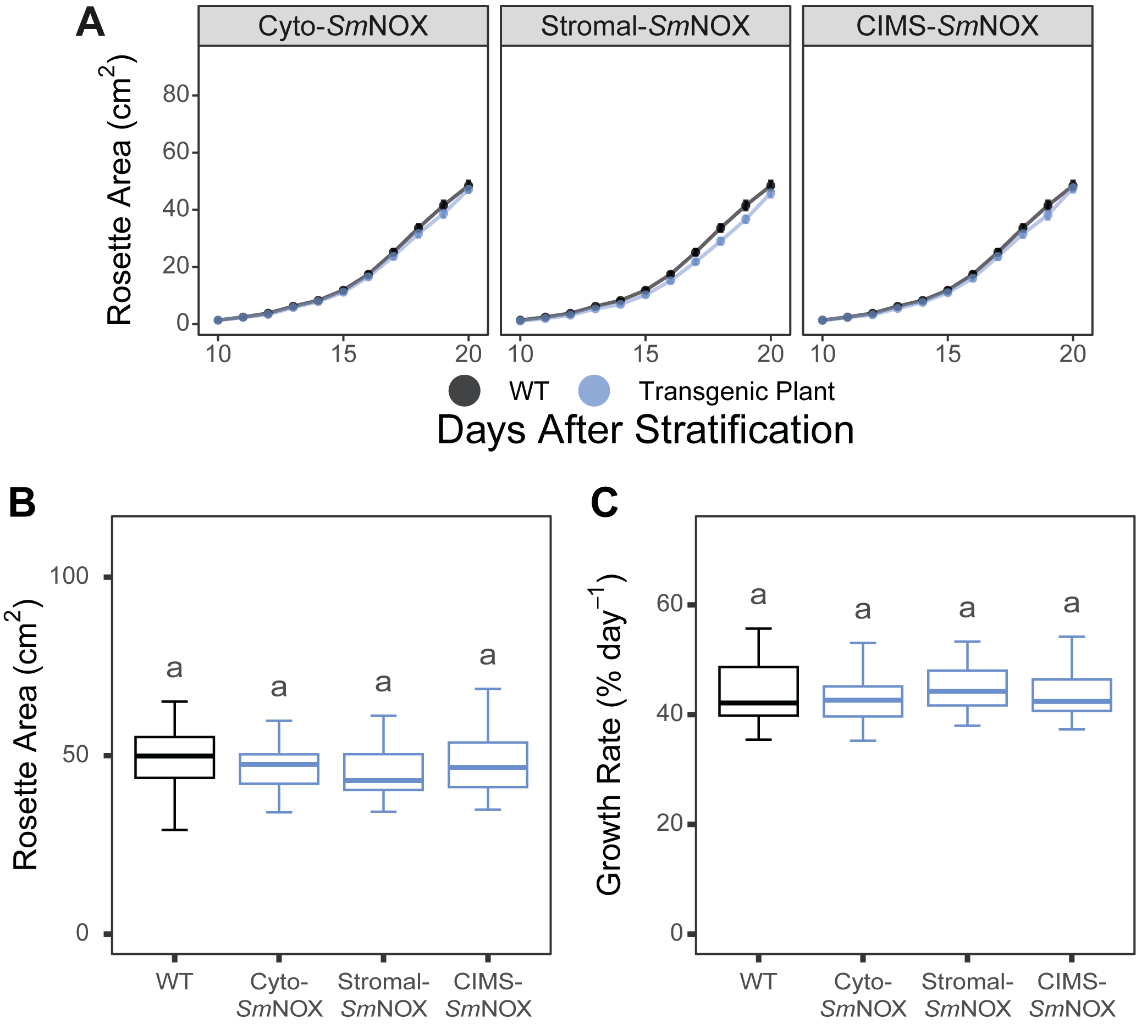

**Figure S9.** The vegetative and reproductive performance of plants under normal watering conditions. (A) The increase in visible rosette area (cm^2^) over time in wild-type (black) and transgenic plants (blue). WT: wild-type. Cyto-*Sm*NOX: cytosolic *Sm*NOX. Stromal-*Sm*NOX: chloroplast stromal *Sm*NOX. CIMS-*Sm*NOX: chloroplast intermembrane space *Sm*NOX. Data represent mean ± 1 S.E. B) Boxplot depicting the maximum visible rosette area at the end of vegetative growth (day = 20). The colour scheme and abbreviations follow that described in (A). C) As in (B), but for the average percentage growth rate of plants computed over the vegetative growth phase (% increase in rosette area day^-1^). All data represent an average across three independent single copy lines (*n* = 10 per individual line for all measurements). Differences between transgenic plants and WT are assessed by Fisher LSD post-hoc analysis following a two-way ANOVA, where letters above each box represent statistically significant differences in mean values (p ≤ 0.05). The raw data can be found in Supplemental File 4.

### Supplemental Tables

#### Supplemental File 1, Table S1

**Table S1.** Predicted localisations of *Sm*NOX (including C-terminal GFP sequence) using target V2.0 software ^14,15^.

|  | **Likelihood** | | | |
| --- | --- | --- | --- | --- |
|  | **Other** | **Mitochondrial TP** | **Chloroplast TP** | **Thylakoid TP** |
| **Cyto-*Sm*NOX** | 0.9948 | 0.0027 | 0 | 0 |

#### Supplemental File 1, Table S2

**Table S2.** Gas exchange parameters derived from light response (*A* – *PPFD*_­_) curves. Values represent mean ± 1 S.E. All data represent an average across three independent single copy lines (*n* = 7 – 8 per individual line). Differences between transgenic plants and WT are assessed by Fisher LSD post-hoc analysis following a two-way ANOVA, where letters above each box represent statistically significant differences in mean values (p ≤ 0.05). Significant differences to WT are also highlighted in bold for ease of visualisation.

|  | **WT** | **Cyto-*Sm*NOX** | **Stromal-*Sm*NOX** | **CIMS-*Sm*NOX** |
| --- | --- | --- | --- | --- |
| ***A* _sat_**  (µmol m^−2^ s^−1^) | 6.2 ± 0.3 ^a^ | 5.3 ± 0.2 ^a^ | 5.8 ± 0.2 ^a^ | 5.2 ± 0.2 ^a^ |
| ***g*_s sat_**  (mol m⁻² s⁻¹) | 0.166 ± 0.008 ^a^ | **0.126 ± 0.006 ^b^** | **0.129 ± 0.005 ^b^** | **0.121 ± 0.006 ^b^** |
| ***iWUE* _sat_**  (µmol mol^-1^ H_2_O) | 41.3 ± 1.4 ^b^ | **46.5 ± 1.6 ^a^** | **48.5 ± 1.0 ^a^** | **48.2 ± 2.2 ^a^** |
| **LCP**  (µmol m^−2^ s^−1^) | 17.7 ± 1.7 ^a^ | 23.3 ± 1.7 ^a^ | 18.5 ± 1.7 ^a^ | 20.5 ± 1.3 ^a^ |
| ***R*_d_**  (µmol m^−2^ s^−1^) | 0.748 ± 0.084 ^a^ | 0.941 ± 0.075 ^a^ | 0.746 ± 0.074 ^a^ | 0.783 ± 0.048 ^a^ |
| ΦJ  (unitless) | 0.043 ± 0.002 ^a^ | 0.041 ± 0.002 ^a^ | 0.040 ± 0.001 ^a^ | 0.040 ± 0.002 ^a^ |
| ***J*_sat_**  (µmol m^−2^ s^−1^) | 59.2 ± 2.1 ^a^ | 57.7 ± 1.9 ^a^ | 57.8 ± 1.6 ^a^ | 54.0 ± 1.5 ^a^ |

***N.B.*** *A*_sat_: Light-saturated assimilation rate. *g*_s sat_: Light-saturated stomatal conductance. *iWUE*_sat_: Light-saturated instantaneous water-use efficiency. LCP: Light compensation point. R_d_: Mitochondrial respiration in the light. ΦJ: Modelled apparent quantum yield computed as the initial slope of the response of J to PPFD. *J*_sat_: Light-saturated electron transport rate.

#### Supplemental File 1, Table S3

**Table S3.** Gas exchange parameters describing the relative coordination between plant photosynthesis and gas exchange derived from light response (*A* – *PPFD*_­_) curves. Values represent mean ± 1 S.E. All data represent an average across three independent single copy lines (*n* = 7 – 8 per individual line). Differences between transgenic plants and WT are assessed by Fisher LSD post-hoc analysis following a two-way ANOVA, where letters above each box represent statistically significant differences in mean values (p ≤ 0.05). Significant differences to WT are also highlighted in bold for ease of visualisation.

|  | **WT** | **Cyto-*Sm*NOX** | **Stromal-*Sm*NOX** | **CIMS-*Sm*NOX** |
| --- | --- | --- | --- | --- |
| ***g_s_* _A95_**  (mol m^−2^ s^−1^) | 0.154 ± 0.008 ^a^ | **0.120 ± 0.006 ^b^** | **0.125 ± 0.005 ^b^** | **0.116 ± 0.006 ^b^** |
| ***g*_s sat_**  (mol m^−2^ s^−1^) | 0.166 ± 0.008 ^a^ | **0.126 ± 0.006 ^b^** | **0.129 ± 0.005 ^b^** | **0.121 ± 0.006 ^b^** |
| Δ ***g_s_***  (mol m^−2^ s^−1^) | 0.012 ± 0.004 ^a^ | **0.005 ± 0.001 ^b^** | **0.004 ± 0.001 ^b^** | **0.005 ± 0.001 ^b^** |
| PPFD **_A95_**  (µmol m^−2^ s^−1^) | 471.4 ± 18.4 ^a^ | 695.2 ± 89.9 ^a^ | 581.0 ± 71.6 ^a^ | 642.9 ± 98.0 ^a^ |
| PPFD ***_g_*_s sat_**  (µmol m^−2^ s^−1^) | 1857.1 ± 245.8 ^a^ | 1523.8 ± 157.9 ^a^ | 1202.4 ± 160.2 ^a^ | 1490.5 ± 175.9 ^a^ |
| Δ PPFD  (µmol m^−2^ s^−1^) | 1385.7 ± 235.5 ^a^ | **828.6 ± 161.3 ^b^** | **621.4 ± 142.8 ^b^** | **847.6 ± 169.7 ^b^** |

***N.B.*** *g*_s_ _A95_: stomatal conductance at the light intensity at which 95% maximum assimilation rate is achieved. *g_s sat_*: Light-saturated stomatal conductance. Δ *g*_s_: The difference in stomatal conductance between the light intensity at which 95% maximum assimilation rate is achieved and the light-saturated stomatal conductance is achieved. PPFD _A95_: Light intensity at which 95% maximum assimilation rate is achieved. PPFD *_gs sat_*: Light intensity at which light-saturated stomatal conductance is achieved. Δ PPFD: The difference in irradiance between the light intensity at which 95% maximum assimilation rate is achieved and the light-saturated stomatal conductance is achieved.

#### Supplemental File 1, Table S4

**Table S4.** Gas exchange parameters derived from CO_2_ response (*A* – *C*_i­_) curves. Values represent mean ± 1 S.E. All data represent an average across three independent single copy lines (*n* = 3 – 5 per individual line). Differences between transgenic plants and WT are assessed by Fisher LSD post-hoc analysis following a two-way ANOVA, where letters above each box represent statistically significant differences in mean values (p ≤ 0.05). Significant differences to WT are also highlighted in bold for ease of visualisation.

|  | **WT** | **Cyto-*Sm*NOX** | **Stromal-*Sm*NOX** | **CIMS-*Sm*NOX** |
| --- | --- | --- | --- | --- |
| ***A* _max_**  (µmol m^−2^ s^−1^) | 13.6 ± 0.9 ^a^ | 12.8 ± 0.9 ^a^ | 14.2 ± 0.5 ^a^ | 11.4 ± 0.6 ^a^ |
| **CCP**  (µmol mol^-1^) | 75.7 ± 1.6 ^a^ | 79.7 ± 2.2 ^a^ | 72.9 ± 1.4 ^a^ | 78.3 ± 1.6 ^a^ |
| ***K*_C_^air^**  (µM) | 1100.4 ± 6.9 ^a^ | 1089.2 ± 7.9 ^a^ | 1092.7 ± 6.9 ^a^ | 1093.8 ± 6.7 ^a^ |
| ***V*_c max_**  (µmol m^−2^ s^−1^) | 37.0 ± 2.6 ^a^ | 37.3 ± 2.4 ^a^ | 39.4 ± 1.9 ^a^ | 31.4 ± 1.5 ^a^ |
| ***V*_c max_ ^25°C^**  (µmol m^−2^ s^−1^) | 21.2 ± 1.6 ^a^ | 21.6 ± 1.4 ^a^ | 22.8 ± 1.1 ^a^ | 18.1 ± 0.9 ^a^ |
| ***J* _max_**  (µmol m^−2^ s^−1^) | 66.1 ± 4.4 ^a^ | 62.8 ± 4.5 ^a^ | 68.7 ± 2.7 ^a^ | 55.0 ± 3.1 ^a^ |
| ***J* _max_ ^25°C^**  (µmol m^−2^ s^−1^) | 54.7 ± 3.7 ^a^ | 52.0 ± 3.8 ^a^ | 56.9 ± 2.2 ^a^ | 45.6 ± 2.5 ^a^ |
| *A*_C_/*A*_J_  (µmol mol^-1^) | 657.6 ± 41.6 ^a^ | 563.3 ± 27.8 ^a^ | 622.7 ± 34.6 ^a^ | 626.4 ± 38.2 ^a^ |

***N.B.*** *A* _max_: CO_2_ saturated assimilation rate. CCP: CO_2_ compensation point. *K*_C_^air^: Modelled Michaelis constant of rubisco for CO_2_ in 21% O_2_ air. *V*_c max_: Modelled maximum carboxylation rate of rubisco at assay temperature of 30°C. *V*_c max_ ^25°C^: Modelled maximum carboxylation rate of rubisco adjusted to 25°C. *J* _max_: Maximum electron transport rate at assay temperature of 30°C. *J* _max_ ^25°C^: Maximum electron transport rate adjusted to 25°C. *A*_C_/*A*_J_: Modelled CO_2_ concentration at transition between rubisco-limited and ribulose-1,5-bisphosphate (RuBP) regeneration-limited photosynthesis.

#### Supplemental File 1, Table S5

**Table S5.** Stomatal kinetic parameters derived from the temporal profiles of plant gas exchange responses to step changes in light intensity between 100 μmol m^−2^ s^−1^ and 1000 μmol m^−2^ s^−1^. Values represent mean ± 1 S.E. All data represent an average across three independent single copy lines (*n* = 3 – 4 per individual line). Differences between transgenic plants and WT are assessed by Fisher LSD post-hoc analysis following a two-way ANOVA, where letters above each box represent statistically significant differences in mean values (p ≤ 0.05). Significant differences to WT are also highlighted in bold for ease of visualisation.

|  | **WT** | **Cyto-*Sm*NOX** | **Stromal-*Sm*NOX** | **CIMS-*Sm*NOX** |
| --- | --- | --- | --- | --- |
| ***λ***  (minutes) | 1.7 ± 0. ^a^ | 1.1 ± 0.1 ^a^ | 1.4 ± 0.2 ^a^ | 1.7 ± 0.3 ^a^ |
| ***k*_i_**  (minutes) | 9.9 ± 0.7 ^a^ | 8.6 ± 0.7 ^a^ | 8.0 ± 0.5 ^a^ | 10.6 ± 0.8 ^a^ |
| ***Sl* _max_**  (mol m^−2^ s^−1^ min^-1^) | 0.065 ± 0.007 ^a^ | 0.101 ± 0.018 ^a^ | 0.096 ± 0.015 ^a^ | 0.046 ± 0.003 ^a^ |
| ***k*_d_** (minutes) | 2.0 ± 0.2 ^a^ | 2.5 ± 0.3 ^a^ | 2.1 ± 0.2 ^a^ | 2.1 ± 0.3 ^a^ |

***N.B.*** λ: Initial lag in response time of stomatal conductance upon step increase in light intensity from 100 μmol m^−2^ s^−1^ and 1000 μmol m^−2^ s^−1^. *k*_i_: Time taken to achieve new steady state stomatal conductance upon step increase in light intensity from 100 μmol m^−2^ s^−1^ and 1000 μmol m^−2^ s^−1^. *Sl* _max_: Maximal rate of stomatal opening upon step increase in light intensity from 100 μmol m^−2^ s^−1^ and 1000 μmol m^−2^ s^−1^ min^-1^. *k*_d_: Time taken to achieve new steady state stomatal conductance upon step decrease in light intensity from 1000 μmol m^−2^ s^−1^ and 100 μmol m^−2^ s^−1^.
