## Supplemental File 2 for "Metabolic engineering of stomatal precursor cells enhances photosynthetic water-use efficiency and vegetative growth under water-deficit conditions in *Arabidopsis thaliana*"

***Sequences***

A list of sequences as used in the present study. Gene sequences which have been commercially synthesized have also been subject to codon optimisation. Thus, the nucleotide sequences described here do not necessarily reflect the native sequences which are found in nature. Different regions of the sequence are highlighted in different colours. Promoter including 5’ untranslated region (UTR): Turquoise. N-terminal fusion protein: Yellow. Translated/untranslated linker sequences: Gray. Fluorescent marker: Bright green. Terminator including 3’ UTR: Pink. Initial start codon (nucleotide sequence) or leading methionine (amino acid sequence): **Bold Text**. Golden Gate compatible flanking regions: Dark Red.

***Nucleotide sequences of synthesized genes***

Full-length nucleotide sequences of each gene (including golden gate flanking regions) as synthesized for cloning (Seq ID 1 – 4).

**SEQ ID 1** – *Nucleotide sequence of the full-length SmNOX with C module overhangs*

cactctgtggtctcaaggt**ATG**AGCAAGATTGTTATCGTTGGAGCCAATCACGCAGGAACCGCTGCAATCAACACTGTGCTGGACAACTATGGATCTGAGAATGAGGTGGTTGTCTTTGACCAGAACTCAAACATTAGCTTCCTGGGATGCGGAATGGCTCTGTGGATCGGTAAGCAGATCTCTGGTCCCCAGGGACTGTTCTATGCAGACAAGGAGTCTCTGGAGGCTAAAGGTGCCAAGATTTACATGGAGAGCCCTGTCACTGCTATCGACTACGATGCTAAACGCGTGACCGCACTGGTCAACGGACAGGAGCACGTCGAATCTTACGAGAAGCTGATCCTGGCCACTGGATCCACTCCCATTCTGCCACCCATCAAGGGTGCAGCAATCAAGGAGGGTAGCCGTGATTTCGAGGCGACCTTGAAGAACCTGCAATTCGTTAAGCTGTATCAGAACGCTGAGGACGTTATTAACAAACTGCAGGACAAGACCCAAAATCTGAACCGTATTGCTGTTGTTGGTGCAGGTTACATTGGTGTCGAACTGGCAGAAGCATTCAAGCGCCTGGGTAAGGAAGTTATTCTGATTGCACGTCACGACACCTGTTTGCGTGGTTACTACGATCAAGATCTATCCGAGATGATGCGTCAGAACTTGGAGGACCACGGCATAGAGCTTGCATTTGGAGAAACTGTTAAAGCAATCGAGGGTGACGGAAAAGTGGAGCGCATCGTCACTGATAAAGCATCTCATGACGTCGACATGGTGATCCTGGCTGTCGGATTCCGCCCCAATACCGCTCTGGGCAACGCTAAGCTGAAAACTTTCCGTAACGGAGCCTTCCTTGTGGATAAGAAACAGGAGACTAGCATTCCAGACGTGTATGCTATTGGTGATTGCGCTACCGTGTACGACAACGCAATCAATGACACTAACTACATCGCGCTGGCATCCAATGCTCTTCGTTCTGGTATTGTAGCCGGACACAACGCTGCTGGACACAAGCTGGAGTCTTTGGGAGTCCAGGGTTCGAATGGTATCTCTATTTTCGGACTGAACATGGTTTCTACCGGTCTGACTCAGGAAAAGGCTAAGCGCTTTGGGTACAATCCTGAAGTCACCGCATTCACTGATTTCCAGAAGGCATCTTTCATCGAGCACGACAACTATCCCGTTACACTGAAGATCGTGTACGACAAAGACTCTCGTCTGGTTCTCGGAGCTCAGATGGCTTCTAAGGAGGATATGTCTATGGGTATCCATATGTTTAGCCTTGCCATTCAGGAGAAGGTTACCATCGAACGTCTGGCACTGCTGGACTATTTCTTTCTTCCCCACTTTAACCAGCCTTACAATTATATGATCAAGGCCGCTCTGAAAGCAAAGTGAgctttgagaccacgaagtg

**SEQ ID 2** – *Nucleotide sequence of Arabidopsis thaliana* *photosystem II subunit QA transit peptide with S module overhangs*

cactctgtggtctcaa**atg**GCTTCTATGGGCGGACTGCACGGGGCCTCCCCCGCAGTCTTGGAGGGAAGCCTGAAGATCAACGGTTCTTCACGTCTGAATGGATCTGGACGCGTGGCTGTTGCACAGCGCTCTCGTCTGGTTGTGCGTGCTCAGCAGTCGGAGGAAACCTCTCGTCGTAGCGTCATTGGTCTTGTCGCAGCGGGTCTGGCAGGTGGTTCTTTCGTTCAAGCTGTGCTGGCTGACGCAATCAGCATTAAAGTTGGACCCCCTCCCGCCggaggttgagaccacgaagtg

**SEQ ID 3** – *Nucleotide sequence of the full-length A. thaliana* *outer envelope protein 9 gene with S module overhangs*

cactctgtggtctcaa**atg**GGTAACGAAACCAAGACTAATGGGGGTCCTGCTAGCATGGCTGGAGGCGGAGGTTTCCGTGCAAAAATGGAGCATTACGTTTACTCTGGTGAAAAGAAGCACGTCCTTGTCGGTATCGGAATCGTTACCATTATTTTTGGAGTGCCCTGGTATCTGATGACTCAGGGATCTAAACACCAGAGCCACCAAGACTATATGGATAAGGCCGACAAGGCACGCAAGGCTCGTCTGTCCTCTTCATCTTCTGCAAACAAAggaggttgagaccacgaagtg

**SEQ ID 4** – *Nucleotide sequence of the full-length A. thaliana* r*ubisco small subunit gene with S module overhangs*

cactctgtggtctcaa**atg**GCTTCTTCTATGCTGTCCAGCGCTACTATGGTGGCATCTCCCGCACAGGCTACCATGGTCGCTCCCTTCAATGGCTTGAAATCTTCCGCTGCCTTCCCCGCAACACGCAAGGCGAACAATGATATCACTAGCATTACCTCTAACGGAGGTCGTGTCAATTGTATGCAGGTGTGGCCTCCCATCGGTAAAAAGAAGTTCGAAACCCTGAGCTACCTGCCTGACCTGACTGACTCTGAGCTGGCAAAGGAGGTTGACTATCTGATCCGTAACAAGTGGATTCCCTGCGTTGAGTTTGAGCTCGAGCATGGGTTCGTTTACCGTGAGCACGGAAACTCTCCCGGATATTACGACGGTCGTTACTGGACCATGTGGAAACTGCCACTTTTCGGTTGCACTGATTCTGCACAGGTCCTTAAGGAAGTTGAGGAATGCAAAAAGGAGTATCCAAACGCATTCATCCGTATCATTGGATTTGACAACACTCGCCAGGTGCAATGCATTTCATTCATCGCCTACAAGCCCCCTTCGTTTACCGGTGGATCTGGTggaggttgagaccacgaagtg

***Nucleotide and translated amino acid sequences of expression cassettes used for stable overexpression***

Full-length sequences of expression cassettes used for stable overexpression of *Sm*NOX (Seq ID 5 – 10). Expression of both untargeted and translationally-fused *Sm*NOX is driven by the *A. thaliana* chlorophyll a/b-binding protein 3 (CAB3) promoter and contain the *Agrobacterium tumefaciens* NOSt terminator.

**SEQ ID 5** – *Nucleotide sequence of the expression cassette of untargeted SmNOX (cyto–SmNOX; cytosolic localisation)*

ggagtctgaagctcgtaacattggctcatacgatatctggataacaatatatcacattgctctgataccatattaaaatcacatccacgcattgaatggactgatcatatatcttacatattagttaaaatccattgtcgtgcgagagattgccttccgtgacatcctctgtggaccaggttcgcttgtcacactagctataattggtgaacacatgataacatatatgtttggagcatagaagagctagcgcctgtgtcattggtgtgtggtatcgaaagaacaagagaatagaacaggagaacatgaatatagctaggtttattcattttcaccccaaagcttagaaattgcgttccaaagagtgggtgaattcatgtgtgagggcaattagtgattgtaaaaataaaattgtgttttgtaaaaaacttttactgtcgaaattatttagggtgatgaaaaaatcagtaaactacgaatgatagcttaaagagtttctatcaaagtgattgaggaatagtttgttgcaaattaaacctctaacaaaatgttttctgttgtggtttttcatctctacaaattttgaattttatgatgaattagaaagatagaatgagttactttagattttaaaaggttgttcaagtttacaaaacagattactagaatcatgattaaaaatttacaagctacatattgtctaaaccaatgatgttgaacataccagatgatagtttttcagtgtttgaacaatcaattggatagtttttatgtttctgcaaaatatgcaaataatcagtgtttttgagtctttgcattttgatttaaaagcaaaaacaactgagtttcaaggttaaattaattacattattcatgagatttatcaggttagtggataaactgacaatggaatcaatgttattgtaaattggtagtgatgttggacttctaatgttactctctatgatgtttcggtcatcaatatcacactatctttacttttatttaaaggaaagatcacacaaataagttatctctattcagaactattaagctgcttccaaaagacttgcaacatgtggactcgaaatgctttggctgcaatgaaaaaatcatagcaaaagctagtggactagagactgccacataagaatagtaaacgttaaaaccaaaatctcaaaaatccaatgagtaaagagatatagattacttcatagataacaaacgttactcgcaattttcctatataatccaaccctacctaaccattttcaatcactctcactcacaagttagtcaccaaaaaaaaaaaaaaacacaaaaagtttca**atg**agcaagattgttatcgttggagccaatcacgcaggaaccgctgcaatcaacactgtgctggacaactatggatctgagaatgaggtggttgtctttgaccagaactcaaacattagcttcctgggatgcggaatggctctgtggatcggtaagcagatctctggtccccagggactgttctatgcagacaaggagtctctggaggctaaaggtgccaagatttacatggagagccctgtcactgctatcgactacgatgctaaacgcgtgaccgcactggtcaacggacaggagcacgtcgaatcttacgagaagctgatcctggccactggatccactcccattctgccacccatcaagggtgcagcaatcaaggagggtagccgtgatttcgaggcgaccttgaagaacctgcaattcgttaagctgtatcagaacgctgaggacgttattaacaaactgcaggacaagacccaaaatctgaaccgtattgctgttgttggtgcaggttacattggtgtcgaactggcagaagcattcaagcgcctgggtaaggaagttattctgattgcacgtcacgacacctgtttgcgtggttactacgatcaagatctatccgagatgatgcgtcagaacttggaggaccacggcatagagcttgcatttggagaaactgttaaagcaatcgagggtgacggaaaagtggagcgcatcgtcactgataaagcatctcatgacgtcgacatggtgatcctggctgtcggattccgccccaataccgctctgggcaacgctaagctgaaaactttccgtaacggagccttccttgtggataagaaacaggagactagcattccagacgtgtatgctattggtgattgcgctaccgtgtacgacaacgcaatcaatgacactaactacatcgcgctggcatccaatgctcttcgttctggtattgtagccggacacaacgctgctggacacaagctggagtctttgggagtccagggttcgaatggtatctctattttcggactgaacatggtttctaccggtctgactcaggaaaaggctaagcgctttgggtacaatcctgaagtcaccgcattcactgatttccagaaggcatctttcatcgagcacgacaactatcccgttacactgaagatcgtgtacgacaaagactctcgtctggttctcggagctcagatggcttctaaggaggatatgtctatgggtatccatatgtttagccttgccattcaggagaaggttaccatcgaacgtctggcactgctggactatttctttcttccccactttaaccagccttacaattatatgatcaaggccgctctgaaagcaaagtgagcttgtcaagcagatcgttcaaacatttggcaataaagtttcttaagattgaatcctgttgccggtcttgcgatgattatcatataatttctgttgaattacgttaagcatgtaataattaacatgtaatgcatgacgttatttatgagatgggtttttatgattagagtcccgcaattatacatttaatacgcgatagaaaacaaaatatagcgcgcaaactaggataaattatcgcgcgcggtgtcatctatgttactagatcga

**SEQ ID 6** – *Nucleotide sequence of the expression cassette of SmNOX translationally fused at the N-termini to the A. thaliana photosystem II subunit QA transit peptide (stromal-SmNOX; chloroplast stromal localisation)*

ggagtctgaagctcgtaacattggctcatacgatatctggataacaatatatcacattgctctgataccatattaaaatcacatccacgcattgaatggactgatcatatatcttacatattagttaaaatccattgtcgtgcgagagattgccttccgtgacatcctctgtggaccaggttcgcttgtcacactagctataattggtgaacacatgataacatatatgtttggagcatagaagagctagcgcctgtgtcattggtgtgtggtatcgaaagaacaagagaatagaacaggagaacatgaatatagctaggtttattcattttcaccccaaagcttagaaattgcgttccaaagagtgggtgaattcatgtgtgagggcaattagtgattgtaaaaataaaattgtgttttgtaaaaaacttttactgtcgaaattatttagggtgatgaaaaaatcagtaaactacgaatgatagcttaaagagtttctatcaaagtgattgaggaatagtttgttgcaaattaaacctctaacaaaatgttttctgttgtggtttttcatctctacaaattttgaattttatgatgaattagaaagatagaatgagttactttagattttaaaaggttgttcaagtttacaaaacagattactagaatcatgattaaaaatttacaagctacatattgtctaaaccaatgatgttgaacataccagatgatagtttttcagtgtttgaacaatcaattggatagtttttatgtttctgcaaaatatgcaaataatcagtgtttttgagtctttgcattttgatttaaaagcaaaaacaactgagtttcaaggttaaattaattacattattcatgagatttatcaggttagtggataaactgacaatggaatcaatgttattgtaaattggtagtgatgttggacttctaatgttactctctatgatgtttcggtcatcaatatcacactatctttacttttatttaaaggaaagatcacacaaataagttatctctattcagaactattaagctgcttccaaaagacttgcaacatgtggactcgaaatgctttggctgcaatgaaaaaatcatagcaaaagctagtggactagagactgccacataagaatagtaaacgttaaaaccaaaatctcaaaaatccaatgagtaaagagatatagattacttcatagataacaaacgttactcgcaattttcctatataatccaaccctacctaaccattttcaatcactctcactcacaagttagtcaccaaaaaaaaaaaaaaacacaaaaagtttca**atg**gcttctatgggcggactgcacggggcctcccccgcagtcttggagggaagcctgaagatcaacggttcttcacgtctgaatggatctggacgcgtggctgttgcacagcgctctcgtctggttgtgcgtgctcagcagtcggaggaaacctctcgtcgtagcgtcattggtcttgtcgcagcgggtctggcaggtggttctttcgttcaagctgtgctggctgacgcaatcagcattaaagttggaccccctcccgccggaggtatgagcaagattgttatcgttggagccaatcacgcaggaaccgctgcaatcaacactgtgctggacaactatggatctgagaatgaggtggttgtctttgaccagaactcaaacattagcttcctgggatgcggaatggctctgtggatcggtaagcagatctctggtccccagggactgttctatgcagacaaggagtctctggaggctaaaggtgccaagatttacatggagagccctgtcactgctatcgactacgatgctaaacgcgtgaccgcactggtcaacggacaggagcacgtcgaatcttacgagaagctgatcctggccactggatccactcccattctgccacccatcaagggtgcagcaatcaaggagggtagccgtgatttcgaggcgaccttgaagaacctgcaattcgttaagctgtatcagaacgctgaggacgttattaacaaactgcaggacaagacccaaaatctgaaccgtattgctgttgttggtgcaggttacattggtgtcgaactggcagaagcattcaagcgcctgggtaaggaagttattctgattgcacgtcacgacacctgtttgcgtggttactacgatcaagatctatccgagatgatgcgtcagaacttggaggaccacggcatagagcttgcatttggagaaactgttaaagcaatcgagggtgacggaaaagtggagcgcatcgtcactgataaagcatctcatgacgtcgacatggtgatcctggctgtcggattccgccccaataccgctctgggcaacgctaagctgaaaactttccgtaacggagccttccttgtggataagaaacaggagactagcattccagacgtgtatgctattggtgattgcgctaccgtgtacgacaacgcaatcaatgacactaactacatcgcgctggcatccaatgctcttcgttctggtattgtagccggacacaacgctgctggacacaagctggagtctttgggagtccagggttcgaatggtatctctattttcggactgaacatggtttctaccggtctgactcaggaaaaggctaagcgctttgggtacaatcctgaagtcaccgcattcactgatttccagaaggcatctttcatcgagcacgacaactatcccgttacactgaagatcgtgtacgacaaagactctcgtctggttctcggagctcagatggcttctaaggaggatatgtctatgggtatccatatgtttagccttgccattcaggagaaggttaccatcgaacgtctggcactgctggactatttctttcttccccactttaaccagccttacaattatatgatcaaggccgctctgaaagcaaagtgagcttgtcaagcagatcgttcaaacatttggcaataaagtttcttaagattgaatcctgttgccggtcttgcgatgattatcatataatttctgttgaattacgttaagcatgtaataattaacatgtaatgcatgacgttatttatgagatgggtttttatgattagagtcccgcaattatacatttaatacgcgatagaaaacaaaatatagcgcgcaaactaggataaattatcgcgcgcggtgtcatctatgttactagatcga

**SEQ ID 7** – *Nucleotide sequence of the expression cassette of SmNOX translationally fused at the N-termini to the full-length A. thaliana outer envelope protein 9 gene (CIMS-SmNOX; chloroplast intermembrane space localisation)*

ggagtctgaagctcgtaacattggctcatacgatatctggataacaatatatcacattgctctgataccatattaaaatcacatccacgcattgaatggactgatcatatatcttacatattagttaaaatccattgtcgtgcgagagattgccttccgtgacatcctctgtggaccaggttcgcttgtcacactagctataattggtgaacacatgataacatatatgtttggagcatagaagagctagcgcctgtgtcattggtgtgtggtatcgaaagaacaagagaatagaacaggagaacatgaatatagctaggtttattcattttcaccccaaagcttagaaattgcgttccaaagagtgggtgaattcatgtgtgagggcaattagtgattgtaaaaataaaattgtgttttgtaaaaaacttttactgtcgaaattatttagggtgatgaaaaaatcagtaaactacgaatgatagcttaaagagtttctatcaaagtgattgaggaatagtttgttgcaaattaaacctctaacaaaatgttttctgttgtggtttttcatctctacaaattttgaattttatgatgaattagaaagatagaatgagttactttagattttaaaaggttgttcaagtttacaaaacagattactagaatcatgattaaaaatttacaagctacatattgtctaaaccaatgatgttgaacataccagatgatagtttttcagtgtttgaacaatcaattggatagtttttatgtttctgcaaaatatgcaaataatcagtgtttttgagtctttgcattttgatttaaaagcaaaaacaactgagtttcaaggttaaattaattacattattcatgagatttatcaggttagtggataaactgacaatggaatcaatgttattgtaaattggtagtgatgttggacttctaatgttactctctatgatgtttcggtcatcaatatcacactatctttacttttatttaaaggaaagatcacacaaataagttatctctattcagaactattaagctgcttccaaaagacttgcaacatgtggactcgaaatgctttggctgcaatgaaaaaatcatagcaaaagctagtggactagagactgccacataagaatagtaaacgttaaaaccaaaatctcaaaaatccaatgagtaaagagatatagattacttcatagataacaaacgttactcgcaattttcctatataatccaaccctacctaaccattttcaatcactctcactcacaagttagtcaccaaaaaaaaaaaaaaacacaaaaagtttca**atg**ggtaacgaaaccaagactaatgggggtcctgctagcatggctggaggcggaggtttccgtgcaaaaatggagcattacgtttactctggtgaaaagaagcacgtccttgtcggtatcggaatcgttaccattatttttggagtgccctggtatctgatgactcagggatctaaacaccagagccaccaagactatatggataaggccgacaaggcacgcaaggctcgtctgtcctcttcatcttctgcaaacaaaggaggtatgagcaagattgttatcgttggagccaatcacgcaggaaccgctgcaatcaacactgtgctggacaactatggatctgagaatgaggtggttgtctttgaccagaactcaaacattagcttcctgggatgcggaatggctctgtggatcggtaagcagatctctggtccccagggactgttctatgcagacaaggagtctctggaggctaaaggtgccaagatttacatggagagccctgtcactgctatcgactacgatgctaaacgcgtgaccgcactggtcaacggacaggagcacgtcgaatcttacgagaagctgatcctggccactggatccactcccattctgccacccatcaagggtgcagcaatcaaggagggtagccgtgatttcgaggcgaccttgaagaacctgcaattcgttaagctgtatcagaacgctgaggacgttattaacaaactgcaggacaagacccaaaatctgaaccgtattgctgttgttggtgcaggttacattggtgtcgaactggcagaagcattcaagcgcctgggtaaggaagttattctgattgcacgtcacgacacctgtttgcgtggttactacgatcaagatctatccgagatgatgcgtcagaacttggaggaccacggcatagagcttgcatttggagaaactgttaaagcaatcgagggtgacggaaaagtggagcgcatcgtcactgataaagcatctcatgacgtcgacatggtgatcctggctgtcggattccgccccaataccgctctgggcaacgctaagctgaaaactttccgtaacggagccttccttgtggataagaaacaggagactagcattccagacgtgtatgctattggtgattgcgctaccgtgtacgacaacgcaatcaatgacactaactacatcgcgctggcatccaatgctcttcgttctggtattgtagccggacacaacgctgctggacacaagctggagtctttgggagtccagggttcgaatggtatctctattttcggactgaacatggtttctaccggtctgactcaggaaaaggctaagcgctttgggtacaatcctgaagtcaccgcattcactgatttccagaaggcatctttcatcgagcacgacaactatcccgttacactgaagatcgtgtacgacaaagactctcgtctggttctcggagctcagatggcttctaaggaggatatgtctatgggtatccatatgtttagccttgccattcaggagaaggttaccatcgaacgtctggcactgctggactatttctttcttccccactttaaccagccttacaattatatgatcaaggccgctctgaaagcaaagtgagcttgtcaagcagatcgttcaaacatttggcaataaagtttcttaagattgaatcctgttgccggtcttgcgatgattatcatataatttctgttgaattacgttaagcatgtaataattaacatgtaatgcatgacgttatttatgagatgggtttttatgattagagtcccgcaattatacatttaatacgcgatagaaaacaaaatatagcgcgcaaactaggataaattatcgcgcgcggtgtcatctatgttactagatcga

**SEQ ID 8** – *Translated amino acid sequence of untargeted SmNOX (cyto-SmNOX; cytosolic localisation)*

**M**SKIVIVGANHAGTAAINTVLDNYGSENEVVVFDQNSNISFLGCGMALWIGKQISGPQGLFYADKESLEAKGAKIYMESPVTAIDYDAKRVTALVNGQEHVESYEKLILATGSTPILPPIKGAAIKEGSRDFEATLKNLQFVKLYQNAEDVINKLQDKTQNLNRIAVVGAGYIGVELAEAFKRLGKEVILIARHDTCLRGYYDQDLSEMMRQNLEDHGIELAFGETVKAIEGDGKVERIVTDKASHDVDMVILAVGFRPNTALGNAKLKTFRNGAFLVDKKQETSIPDVYAIGDCATVYDNAINDTNYIALASNALRSGIVAGHNAAGHKLESLGVQGSNGISIFGLNMVSTGLTQEKAKRFGYNPEVTAFTDFQKASFIEHDNYPVTLKIVYDKDSRLVLGAQMASKEDMSMGIHMFSLAIQEKVTIERLALLDYFFLPHFNQPYNYMIKAALKAK*

**SEQ ID 9** – *Translated amino acid sequence of SmNOX translationally fused at the N-termini to the A. thaliana photosystem II subunit QA transit peptide (stromal-SmNOX; chloroplast stromal localisation)*

**M**ASMGGLHGASPAVLEGSLKINGSSRLNGSGRVAVAQRSRLVVRAQQSEETSRRSVIGLVAAGLAGGSFVQAVLADAISIKVGPPPAGGMSKIVIVGANHAGTAAINTVLDNYGSENEVVVFDQNSNISFLGCGMALWIGKQISGPQGLFYADKESLEAKGAKIYMESPVTAIDYDAKRVTALVNGQEHVESYEKLILATGSTPILPPIKGAAIKEGSRDFEATLKNLQFVKLYQNAEDVINKLQDKTQNLNRIAVVGAGYIGVELAEAFKRLGKEVILIARHDTCLRGYYDQDLSEMMRQNLEDHGIELAFGETVKAIEGDGKVERIVTDKASHDVDMVILAVGFRPNTALGNAKLKTFRNGAFLVDKKQETSIPDVYAIGDCATVYDNAINDTNYIALASNALRSGIVAGHNAAGHKLESLGVQGSNGISIFGLNMVSTGLTQEKAKRFGYNPEVTAFTDFQKASFIEHDNYPVTLKIVYDKDSRLVLGAQMASKEDMSMGIHMFSLAIQEKVTIERLALLDYFFLPHFNQPYNYMIKAALKAK*

**SEQ ID 10** – *Translated amino acid sequence of SmNOX translationally fused at the N-termini to the full-length A. thaliana outer envelope protein 9 gene (CIMS-SmNOX; chloroplast intermembrane space localisation)*

**M**GNETKTNGGPASMAGGGGFRAKMEHYVYSGEKKHVLVGIGIVTIIFGVPWYLMTQGSKHQSHQDYMDKADKARKARLSSSSSANKGGMSKIVIVGANHAGTAAINTVLDNYGSENEVVVFDQNSNISFLGCGMALWIGKQISGPQGLFYADKESLEAKGAKIYMESPVTAIDYDAKRVTALVNGQEHVESYEKLILATGSTPILPPIKGAAIKEGSRDFEATLKNLQFVKLYQNAEDVINKLQDKTQNLNRIAVVGAGYIGVELAEAFKRLGKEVILIARHDTCLRGYYDQDLSEMMRQNLEDHGIELAFGETVKAIEGDGKVERIVTDKASHDVDMVILAVGFRPNTALGNAKLKTFRNGAFLVDKKQETSIPDVYAIGDCATVYDNAINDTNYIALASNALRSGIVAGHNAAGHKLESLGVQGSNGISIFGLNMVSTGLTQEKAKRFGYNPEVTAFTDFQKASFIEHDNYPVTLKIVYDKDSRLVLGAQMASKEDMSMGIHMFSLAIQEKVTIERLALLDYFFLPHFNQPYNYMIKAALKAK*

***Nucleotide and translated amino acid sequences of the expression cassette containing the selectable marker used for stable plant transformation***

Full-length sequence of the expression cassette encoding a seed coat specific fluorescent selectable marker used for stable plant transformation (Seq ID 11 – 13). Expression of translationally-fused TagRFP is driven by the native *A. thaliana* oleosin1 promoter (OLEp) and terminator (OLEt), respectively.

**SEQ ID 11** – *Nucleotide sequence of the expression cassette of the TagRFP fluorescent selectable marker translationally fused at the N-termini to the full-length A. thaliana oleosin1 gene (including an intron) (TagRFP; seed-coat specific fluorescence)*

aatgtcgcggaacaaattttaaaactaaatcctaaatttttctaattttgttgccaatagtggatatgtgggccgtatagaaggaatctattgaaggcccaaacccatactgacgagcccaaaggttcgttttgcgttttatgtttcggttcgatgccaacgccacattctgagctaggcaaaaaacaaacgtgtctttgaatagactcctctcgttaacacatgcagcggctgcatggtgacgccattaacacgtggcctacaattgcatgatgtctccattgacacgtgacttctcgtctcctttcttaatatatctaacaaacactcctacctcttccaaaatatatacacatctttttgatcaatctctcattcaaaatctcattctctctagtaaacaagaacaaaaaaatggcggatacagctagaggaacccatcacgatatcatcggcagagatcagtacccgatgatgggccgagatcgtgaccagtaccagatgtccggacgaggatctgactactccaagtctaggcagattgctaaagctgcaactgctgtcacagctggtggttccctccttgttctctccagccttacccttgttggaactgtcatagctttgactgttgcaacacctctgctcgttatcttcagcccaatccttgtcccggctctcatcacagttgcactcctcatcaccggttttctttcctctggagggtttggcattgccgctataaccgttttctcttggatttacaagtaagcacacatttatcatcttacttcataattttgtgcaatatgtgcatgcatgtgttgagccagtagctttggatcaatttttttggtcgaataacaaatgtaacaataagaaattgcaaattctagggaacatttggttaactaaatacgaaatttgacctagctagcttgaatgtgtctgtgtatatcatctatataggtaaaatgcttggtatgatacctattgattgtgaataggtacgcaacgggagagcacccacagggatcagacaagttggacagtgcaaggatgaagttgggaagcaaagctcaggatctgaaagacagagctcagtactacggacagcaacatactggtggggaacatgaccgtgaccgtactcgtggtggccagcacactactatgagcgagctgattaaggagaacatgcacatgaagctgtacatggagggcaccgtgaacaaccaccacttcaagtgcacatccgagggcgaaggcaagccctacgagggcacccagaccatgagaatcaaggtggtcgagggcggccctctccccttcgccttcgacatcctggctaccagcttcatgtacggcagcagaaccttcatcaaccacacccagggcatccccgacttctttaagcagtccttccctgagggcttcacatgggagagagtcaccacatacgaagatgggggcgtgctgaccgctacccaggacaccagcctccaggacggctgcctcatctacaacgtcaagatcagaggggtgaacttcccatccaacggccctgtgatgcagaagaaaacactcggctgggaggccaacaccgagatgctgtaccccgctgacggcggcctggaaggcagaagcgacatggccctgaagctcgtgggcgggggccacctgatctgcaacttcaagaccacatacagatccaagaaacccgctaagaacctcaagatgcccggcgtctactatgtggaccacagactggaaagaatcaaggaggccgacaaagaaacctacgtcgagcagcacgaggtggctgtggccagatactgcgacctccctagcaaactggggcacaagtgagcttaccccactgatgtcatcgtcatagtccaataactccaatgtcggggagttagtttatgaggaataaagtgtttagaatttgatcagggggagataataaaagccgagtttgaatctttttgttataagtaatgtttatgtgtgtttctatatgttgtcaaatggtaccatgttttttttcctctctttttgtaacttgcaagtgttgtgttgtactttatttggcttctttgtaagttggtaacggtggtctatatatggaaaaggtcttgttttgttaaacttatgttagttaactggattcgtctttaaccacaaaaagttttcaataagctacaaatttagacacgcaagccgatgcagtcattagtacatatatttattgcaagtgattacatggcaacccaaacttcaaaaacagtaggttgctccatttagt

**SEQ ID 12** – *Nucleotide sequence of the expression cassette of the TagRFP fluorescent selectable marker translationally fused at the N-termini to the full-length A. thaliana oleosin1 gene (intron removed) (TagRFP; seed-coat specific fluorescence)*

Aatgtcgcggaacaaattttaaaactaaatcctaaatttttctaattttgttgccaatagtggatatgtgggccgtatagaaggaatctattgaaggcccaaacccatactgacgagcccaaaggttcgttttgcgttttatgtttcggttcgatgccaacgccacattctgagctaggcaaaaaacaaacgtgtctttgaatagactcctctcgttaacacatgcagcggctgcatggtgacgccattaacacgtggcctacaattgcatgatgtctccattgacacgtgacttctcgtctcctttcttaatatatctaacaaacactcctacctcttccaaaatatatacacatctttttgatcaatctctcattcaaaatctcattctctctagtaaacaagaacaaaaaaatggcggatacagctagaggaacccatcacgatatcatcggcagagatcagtacccgatgatgggccgagatcgtgaccagtaccagatgtccggacgaggatctgactactccaagtctaggcagattgctaaagctgcaactgctgtcacagctggtggttccctccttgttctctccagccttacccttgttggaactgtcatagctttgactgttgcaacacctctgctcgttatcttcagcccaatccttgtcccggctctcatcacagttgcactcctcatcaccggttttctttcctctggagggtttggcattgccgctataaccgttttctcttggatttacaagtacgcaacgggagagcacccacagggatcagacaagttggacagtgcaaggatgaagttgggaagcaaagctcaggatctgaaagacagagctcagtactacggacagcaacatactggtggggaacatgaccgtgaccgtactcgtggtggccagcacactactatgagcgagctgattaaggagaacatgcacatgaagctgtacatggagggcaccgtgaacaaccaccacttcaagtgcacatccgagggcgaaggcaagccctacgagggcacccagaccatgagaatcaaggtggtcgagggcggccctctccccttcgccttcgacatcctggctaccagcttcatgtacggcagcagaaccttcatcaaccacacccagggcatccccgacttctttaagcagtccttccctgagggcttcacatgggagagagtcaccacatacgaagatgggggcgtgctgaccgctacccaggacaccagcctccaggacggctgcctcatctacaacgtcaagatcagaggggtgaacttcccatccaacggccctgtgatgcagaagaaaacactcggctgggaggccaacaccgagatgctgtaccccgctgacggcggcctggaaggcagaagcgacatggccctgaagctcgtgggcgggggccacctgatctgcaacttcaagaccacatacagatccaagaaacccgctaagaacctcaagatgcccggcgtctactatgtggaccacagactggaaagaatcaaggaggccgacaaagaaacctacgtcgagcagcacgaggtggctgtggccagatactgcgacctccctagcaaactggggcacaagtgagcttaccccactgatgtcatcgtcatagtccaataactccaatgtcggggagttagtttatgaggaataaagtgtttagaatttgatcagggggagataataaaagccgagtttgaatctttttgttataagtaatgtttatgtgtgtttctatatgttgtcaaatggtaccatgttttttttcctctctttttgtaacttgcaagtgttgtgttgtactttatttggcttctttgtaagttggtaacggtggtctatatatggaaaaggtcttgttttgttaaacttatgttagttaactggattcgtctttaaccacaaaaagttttcaataagctacaaatttagacacgcaagccgatgcagtcattagtacatatatttattgcaagtgattacatggcaacccaaacttcaaaaacagtaggttgctccatttagt

**SEQ ID 13** – *Translated amino acid sequence of the expression cassette of the TagRFP fluorescent selectable marker translationally fused at the N-termini to the full-length A. thaliana oleosin1 gene (TagRFP; seed-coat specific fluorescence)*

**M**ADTARGTHHDIIGRDQYPMMGRDRDQYQMSGRGSDYSKSRQIAKAATAVTAGGSLLVLSSLTLVGTVIALTVATPLLVIFSPILVPALITVALLITGFLSSGGFGIAAITVFSWIYKYATGEHPQGSDKLDSARMKLGSKAQDLKDRAQYYGQQHTGGEHDRDRTRGGQHTTMSELIKENMHMKLYMEGTVNNHHFKCTSEGEGKPYEGTQTMRIKVVEGGPLPFAFDILATSFMYGSRTFINHTQGIPDFFKQSFPEGFTWERVTTYEDGGVLTATQDTSLQDGCLIYNVKIRGVNFPSNGPVMQKKTLGWEANTEMLYPADGGLEGRSDMALKLVGGGHLICNFKTTYRSKKPAKNLKMPGVYYVDHRLERIKEADKETYVEQHEVAVARYCDLPSKLGHK*

***Nucleotide and translated amino acid sequences of expression cassettes used for localisation experiments***

Full-length sequences of expression cassettes used for transient expression of *Sm*NOX in protoplasts (Seq ID 14 – 19). Expression of both untargeted and translationally-fused *Sm*NOX is driven by the cauliflower mosaic virus 35S promoter (CaMV35S) promoter and contain the *A. tumefaciens* nopaline synthase terminator (NOSt). All sequences contain a C-terminal GFP.

**SEQ ID 14** – *Nucleotide sequence of the expression cassette of untargeted SmNOX translationally fused to GFP at the C-terminus (cyto-SmNOX; cytosolic localisation)*

tcaacatggtggagcacgacactctggtctactccaaaaatgtcaaagatacagtctcagaagatcaaagggctattgagacttttcaacaaaggataatttcgggaaacctcctcggattccattgcccagctatctgtcacttcatcgaaaggacagtagaaaaggaaggtggctcctacaaatgccatcattgcgataaaggaaaggctatcattcaagatctctctgccgacagtggtcccaaagatggacccccacccacgaggagcatcgtggaaaaagaagaggttccaaccacgtctacaaagcaagtggattgatgtgacatctccactgacgtaagggatgacgcacaatcccactatccttcgcaagacccttcctctatataaggaagttcatttcatttggagaggacacgca**atg**agcaagattgttatcgttggagccaatcacgcaggaaccgctgcaatcaacactgtgctggacaactatggatctgagaatgaggtggttgtctttgaccagaactcaaacattagcttcctgggatgcggaatggctctgtggatcggtaagcagatctctggtccccagggactgttctatgcagacaaggagtctctggaggctaaaggtgccaagatttacatggagagccctgtcactgctatcgactacgatgctaaacgcgtgaccgcactggtcaacggacaggagcacgtcgaatcttacgagaagctgatcctggccactggatccactcccattctgccacccatcaagggtgcagcaatcaaggagggtagccgtgatttcgaggcgaccttgaagaacctgcaattcgttaagctgtatcagaacgctgaggacgttattaacaaactgcaggacaagacccaaaatctgaaccgtattgctgttgttggtgcaggttacattggtgtcgaactggcagaagcattcaagcgcctgggtaaggaagttattctgattgcacgtcacgacacctgtttgcgtggttactacgatcaagatctatccgagatgatgcgtcagaacttggaggaccacggcatagagcttgcatttggagaaactgttaaagcaatcgagggtgacggaaaagtggagcgcatcgtcactgataaagcatctcatgacgtcgacatggtgatcctggctgtcggattccgccccaataccgctctgggcaacgctaagctgaaaactttccgtaacggagccttccttgtggataagaaacaggagactagcattccagacgtgtatgctattggtgattgcgctaccgtgtacgacaacgcaatcaatgacactaactacatcgcgctggcatccaatgctcttcgttctggtattgtagccggacacaacgctgctggacacaagctggagtctttgggagtccagggttcgaatggtatctctattttcggactgaacatggtttctaccggtctgactcaggaaaaggctaagcgctttgggtacaatcctgaagtcaccgcattcactgatttccagaaggcatctttcatcgagcacgacaactatcccgttacactgaagatcgtgtacgacaaagactctcgtctggttctcggagctcagatggcttctaaggaggatatgtctatgggtatccatatgtttagccttgccattcaggagaaggttaccatcgaacgtctggcactgctggactatttctttcttccccactttaaccagccttacaattatatgatcaaggccgctctgaaagcaaagggtggttcggctgtgagcaagggcgaggagctgttcaccggggtggtgcccatcctggtcgagctggacggcgacgtaaacggccacaagttcagcgtgtccggcgagggcgagggcgatgccacctacggcaagctgaccctgaagttcatctgcaccaccggcaagctgcccgtgccctggcccaccctcgtgaccaccctgacctacggcgtgcagtgcttcagccgctaccccgaccacatgaagcagcacgacttcttcaagtccgccatgcccgaaggctacgtccaggagcgcaccatcttcttcaaggacgacggcaactacaagacccgcgccgaggtgaagttcgagggcgacaccctggtgaaccgcatcgagctgaagggcatcgacttcaaggaggacggcaacatcctggggcacaagctggagtacaactacaacagccacaacgtctatatcatggccgacaagcagaagaacggcatcaaggtgaacttcaagatccgccacaacatcgaggacggcagcgtgcagctcgccgaccactaccagcagaacacccccatcggcgacggccccgtgctgctgcccgacaaccactacctgagtactcagtccaagctgagcaaagaccccaacgagaagcgcgatcacatggtcctgctggagttcgtgaccgccgccgggatcactctcggcatggacgagctgtacaaatgagcttgtcaagcagatcgttcaaacatttggcaataaagtttcttaagattgaatcctgttgccggtcttgcgatgattatcatataatttctgttgaattacgttaagcatgtaataattaacatgtaatgcatgacgttatttatgagatgggtttttatgattagagtcccgcaattatacatttaatacgcgatagaaaacaaaatatagcgcgcaaactaggataaattatcgcgcgcggtgtcatctatgttactagatcga

**SEQ ID 15** – *Nucleotide sequence of the expression cassette of SmNOX translationally fused to GFP at the C-terminus and translationally fused at the N-termini to the A. thaliana* *photosystem II subunit QA transit peptide (stromal-SmNOX; chloroplast stromal localisation)*

tcaacatggtggagcacgacactctggtctactccaaaaatgtcaaagatacagtctcagaagatcaaagggctattgagacttttcaacaaaggataatttcgggaaacctcctcggattccattgcccagctatctgtcacttcatcgaaaggacagtagaaaaggaaggtggctcctacaaatgccatcattgcgataaaggaaaggctatcattcaagatctctctgccgacagtggtcccaaagatggacccccacccacgaggagcatcgtggaaaaagaagaggttccaaccacgtctacaaagcaagtggattgatgtgacatctccactgacgtaagggatgacgcacaatcccactatccttcgcaagacccttcctctatataaggaagttcatttcatttggagaggacacgcaatggcttctatgggcggactgcacggggcctcccccgcagtcttggagggaagcctgaagatcaacggttcttcacgtctgaatggatctggacgcgtggctgttgcacagcgctctcgtctggttgtgcgtgctcagcagtcggaggaaacctctcgtcgtagcgtcattggtcttgtcgcagcgggtctggcaggtggttctttcgttcaagctgtgctggctgacgcaatcagcattaaagttggaccccctcccgccggaggt**atg**agcaagattgttatcgttggagccaatcacgcaggaaccgctgcaatcaacactgtgctggacaactatggatctgagaatgaggtggttgtctttgaccagaactcaaacattagcttcctgggatgcggaatggctctgtggatcggtaagcagatctctggtccccagggactgttctatgcagacaaggagtctctggaggctaaaggtgccaagatttacatggagagccctgtcactgctatcgactacgatgctaaacgcgtgaccgcactggtcaacggacaggagcacgtcgaatcttacgagaagctgatcctggccactggatccactcccattctgccacccatcaagggtgcagcaatcaaggagggtagccgtgatttcgaggcgaccttgaagaacctgcaattcgttaagctgtatcagaacgctgaggacgttattaacaaactgcaggacaagacccaaaatctgaaccgtattgctgttgttggtgcaggttacattggtgtcgaactggcagaagcattcaagcgcctgggtaaggaagttattctgattgcacgtcacgacacctgtttgcgtggttactacgatcaagatctatccgagatgatgcgtcagaacttggaggaccacggcatagagcttgcatttggagaaactgttaaagcaatcgagggtgacggaaaagtggagcgcatcgtcactgataaagcatctcatgacgtcgacatggtgatcctggctgtcggattccgccccaataccgctctgggcaacgctaagctgaaaactttccgtaacggagccttccttgtggataagaaacaggagactagcattccagacgtgtatgctattggtgattgcgctaccgtgtacgacaacgcaatcaatgacactaactacatcgcgctggcatccaatgctcttcgttctggtattgtagccggacacaacgctgctggacacaagctggagtctttgggagtccagggttcgaatggtatctctattttcggactgaacatggtttctaccggtctgactcaggaaaaggctaagcgctttgggtacaatcctgaagtcaccgcattcactgatttccagaaggcatctttcatcgagcacgacaactatcccgttacactgaagatcgtgtacgacaaagactctcgtctggttctcggagctcagatggcttctaaggaggatatgtctatgggtatccatatgtttagccttgccattcaggagaaggttaccatcgaacgtctggcactgctggactatttctttcttccccactttaaccagccttacaattatatgatcaaggccgctctgaaagcaaagggtggttcggctgtgagcaagggcgaggagctgttcaccggggtggtgcccatcctggtcgagctggacggcgacgtaaacggccacaagttcagcgtgtccggcgagggcgagggcgatgccacctacggcaagctgaccctgaagttcatctgcaccaccggcaagctgcccgtgccctggcccaccctcgtgaccaccctgacctacggcgtgcagtgcttcagccgctaccccgaccacatgaagcagcacgacttcttcaagtccgccatgcccgaaggctacgtccaggagcgcaccatcttcttcaaggacgacggcaactacaagacccgcgccgaggtgaagttcgagggcgacaccctggtgaaccgcatcgagctgaagggcatcgacttcaaggaggacggcaacatcctggggcacaagctggagtacaactacaacagccacaacgtctatatcatggccgacaagcagaagaacggcatcaaggtgaacttcaagatccgccacaacatcgaggacggcagcgtgcagctcgccgaccactaccagcagaacacccccatcggcgacggccccgtgctgctgcccgacaaccactacctgagtactcagtccaagctgagcaaagaccccaacgagaagcgcgatcacatggtcctgctggagttcgtgaccgccgccgggatcactctcggcatggacgagctgtacaaatgagcttgtcaagcagatcgttcaaacatttggcaataaagtttcttaagattgaatcctgttgccggtcttgcgatgattatcatataatttctgttgaattacgttaagcatgtaataattaacatgtaatgcatgacgttatttatgagatgggtttttatgattagagtcccgcaattatacatttaatacgcgatagaaaacaaaatatagcgcgcaaactaggataaattatcgcgcgcggtgtcatctatgttactagatcga

**SEQ ID 16** – *Nucleotide sequence of the expression cassette of SmNOX translationally fused to GFP at the C-terminus and translationally fused at the N-termini to the full-length A. thaliana outer envelope protein 9 gene (CIMS-SmNOX; chloroplast intermembrane space localisation)*

tcaacatggtggagcacgacactctggtctactccaaaaatgtcaaagatacagtctcagaagatcaaagggctattgagacttttcaacaaaggataatttcgggaaacctcctcggattccattgcccagctatctgtcacttcatcgaaaggacagtagaaaaggaaggtggctcctacaaatgccatcattgcgataaaggaaaggctatcattcaagatctctctgccgacagtggtcccaaagatggacccccacccacgaggagcatcgtggaaaaagaagaggttccaaccacgtctacaaagcaagtggattgatgtgacatctccactgacgtaagggatgacgcacaatcccactatccttcgcaagacccttcctctatataaggaagttcatttcatttggagaggacacgcaatgggtaacgaaaccaagactaatgggggtcctgctagcatggctggaggcggaggtttccgtgcaaaaatggagcattacgtttactctggtgaaaagaagcacgtccttgtcggtatcggaatcgttaccattatttttggagtgccctggtatctgatgactcagggatctaaacaccagagccaccaagactatatggataaggccgacaaggcacgcaaggctcgtctgtcctcttcatcttctgcaaacaaaggaggt**atg**agcaagattgttatcgttggagccaatcacgcaggaaccgctgcaatcaacactgtgctggacaactatggatctgagaatgaggtggttgtctttgaccagaactcaaacattagcttcctgggatgcggaatggctctgtggatcggtaagcagatctctggtccccagggactgttctatgcagacaaggagtctctggaggctaaaggtgccaagatttacatggagagccctgtcactgctatcgactacgatgctaaacgcgtgaccgcactggtcaacggacaggagcacgtcgaatcttacgagaagctgatcctggccactggatccactcccattctgccacccatcaagggtgcagcaatcaaggagggtagccgtgatttcgaggcgaccttgaagaacctgcaattcgttaagctgtatcagaacgctgaggacgttattaacaaactgcaggacaagacccaaaatctgaaccgtattgctgttgttggtgcaggttacattggtgtcgaactggcagaagcattcaagcgcctgggtaaggaagttattctgattgcacgtcacgacacctgtttgcgtggttactacgatcaagatctatccgagatgatgcgtcagaacttggaggaccacggcatagagcttgcatttggagaaactgttaaagcaatcgagggtgacggaaaagtggagcgcatcgtcactgataaagcatctcatgacgtcgacatggtgatcctggctgtcggattccgccccaataccgctctgggcaacgctaagctgaaaactttccgtaacggagccttccttgtggataagaaacaggagactagcattccagacgtgtatgctattggtgattgcgctaccgtgtacgacaacgcaatcaatgacactaactacatcgcgctggcatccaatgctcttcgttctggtattgtagccggacacaacgctgctggacacaagctggagtctttgggagtccagggttcgaatggtatctctattttcggactgaacatggtttctaccggtctgactcaggaaaaggctaagcgctttgggtacaatcctgaagtcaccgcattcactgatttccagaaggcatctttcatcgagcacgacaactatcccgttacactgaagatcgtgtacgacaaagactctcgtctggttctcggagctcagatggcttctaaggaggatatgtctatgggtatccatatgtttagccttgccattcaggagaaggttaccatcgaacgtctggcactgctggactatttctttcttccccactttaaccagccttacaattatatgatcaaggccgctctgaaagcaaagggtggttcggctgtgagcaagggcgaggagctgttcaccggggtggtgcccatcctggtcgagctggacggcgacgtaaacggccacaagttcagcgtgtccggcgagggcgagggcgatgccacctacggcaagctgaccctgaagttcatctgcaccaccggcaagctgcccgtgccctggcccaccctcgtgaccaccctgacctacggcgtgcagtgcttcagccgctaccccgaccacatgaagcagcacgacttcttcaagtccgccatgcccgaaggctacgtccaggagcgcaccatcttcttcaaggacgacggcaactacaagacccgcgccgaggtgaagttcgagggcgacaccctggtgaaccgcatcgagctgaagggcatcgacttcaaggaggacggcaacatcctggggcacaagctggagtacaactacaacagccacaacgtctatatcatggccgacaagcagaagaacggcatcaaggtgaacttcaagatccgccacaacatcgaggacggcagcgtgcagctcgccgaccactaccagcagaacacccccatcggcgacggccccgtgctgctgcccgacaaccactacctgagtactcagtccaagctgagcaaagaccccaacgagaagcgcgatcacatggtcctgctggagttcgtgaccgccgccgggatcactctcggcatggacgagctgtacaaatgagcttgtcaagcagatcgttcaaacatttggcaataaagtttcttaagattgaatcctgttgccggtcttgcgatgattatcatataatttctgttgaattacgttaagcatgtaataattaacatgtaatgcatgacgttatttatgagatgggtttttatgattagagtcccgcaattatacatttaatacgcgatagaaaacaaaatatagcgcgcaaactaggataaattatcgcgcgcggtgtcatctatgttactagatcga

**SEQ ID 17** – *Translated amino acid sequence of untargeted SmNOX translationally fused to GFP at the C-terminus (cyto-SmNOX; cytosolic localisation)*

**M**SKIVIVGANHAGTAAINTVLDNYGSENEVVVFDQNSNISFLGCGMALWIGKQISGPQGLFYADKESLEAKGAKIYMESPVTAIDYDAKRVTALVNGQEHVESYEKLILATGSTPILPPIKGAAIKEGSRDFEATLKNLQFVKLYQNAEDVINKLQDKTQNLNRIAVVGAGYIGVELAEAFKRLGKEVILIARHDTCLRGYYDQDLSEMMRQNLEDHGIELAFGETVKAIEGDGKVERIVTDKASHDVDMVILAVGFRPNTALGNAKLKTFRNGAFLVDKKQETSIPDVYAIGDCATVYDNAINDTNYIALASNALRSGIVAGHNAAGHKLESLGVQGSNGISIFGLNMVSTGLTQEKAKRFGYNPEVTAFTDFQKASFIEHDNYPVTLKIVYDKDSRLVLGAQMASKEDMSMGIHMFSLAIQEKVTIERLALLDYFFLPHFNQPYNYMIKAALKAKGGSAVSKGEELFTGVVPILVELDGDVNGHKFSVSGEGEGDATYGKLTLKFICTTGKLPVPWPTLVTTLTYGVQCFSRYPDHMKQHDFFKSAMPEGYVQERTIFFKDDGNYKTRAEVKFEGDTLVNRIELKGIDFKEDGNILGHKLEYNYNSHNVYIMADKQKNGIKVNFKIRHNIEDGSVQLADHYQQNTPIGDGPVLLPDNHYLSTQSKLSKDPNEKRDHMVLLEFVTAAGITLGMDELYK*

**SEQ ID 18** – *Translated amino acid sequence of SmNOX translationally fused to GFP at the C-terminus and translationally fused at the N-termini to the A. thaliana from the photosystem II subunit QA transit peptide (stromal-SmNOX; chloroplast stromal localisation)*

**M**ASMGGLHGASPAVLEGSLKINGSSRLNGSGRVAVAQRSRLVVRAQQSEETSRRSVIGLVAAGLAGGSFVQAVLADAISIKVGPPPAGGMSKIVIVGANHAGTAAINTVLDNYGSENEVVVFDQNSNISFLGCGMALWIGKQISGPQGLFYADKESLEAKGAKIYMESPVTAIDYDAKRVTALVNGQEHVESYEKLILATGSTPILPPIKGAAIKEGSRDFEATLKNLQFVKLYQNAEDVINKLQDKTQNLNRIAVVGAGYIGVELAEAFKRLGKEVILIARHDTCLRGYYDQDLSEMMRQNLEDHGIELAFGETVKAIEGDGKVERIVTDKASHDVDMVILAVGFRPNTALGNAKLKTFRNGAFLVDKKQETSIPDVYAIGDCATVYDNAINDTNYIALASNALRSGIVAGHNAAGHKLESLGVQGSNGISIFGLNMVSTGLTQEKAKRFGYNPEVTAFTDFQKASFIEHDNYPVTLKIVYDKDSRLVLGAQMASKEDMSMGIHMFSLAIQEKVTIERLALLDYFFLPHFNQPYNYMIKAALKAKGGSAVSKGEELFTGVVPILVELDGDVNGHKFSVSGEGEGDATYGKLTLKFICTTGKLPVPWPTLVTTLTYGVQCFSRYPDHMKQHDFFKSAMPEGYVQERTIFFKDDGNYKTRAEVKFEGDTLVNRIELKGIDFKEDGNILGHKLEYNYNSHNVYIMADKQKNGIKVNFKIRHNIEDGSVQLADHYQQNTPIGDGPVLLPDNHYLSTQSKLSKDPNEKRDHMVLLEFVTAAGITLGMDELYK*

**SEQ ID 19** – *Translated amino acid sequence of SmNOX translationally fused to GFP at the C-terminus and translationally fused at the N-termini to the full-length A. thaliana outer envelope protein 9 gene (CIMS-SmNOX; chloroplast intermembrane space localisation)*

**M**GNETKTNGGPASMAGGGGFRAKMEHYVYSGEKKHVLVGIGIVTIIFGVPWYLMTQGSKHQSHQDYMDKADKARKARLSSSSSANKGGMSKIVIVGANHAGTAAINTVLDNYGSENEVVVFDQNSNISFLGCGMALWIGKQISGPQGLFYADKESLEAKGAKIYMESPVTAIDYDAKRVTALVNGQEHVESYEKLILATGSTPILPPIKGAAIKEGSRDFEATLKNLQFVKLYQNAEDVINKLQDKTQNLNRIAVVGAGYIGVELAEAFKRLGKEVILIARHDTCLRGYYDQDLSEMMRQNLEDHGIELAFGETVKAIEGDGKVERIVTDKASHDVDMVILAVGFRPNTALGNAKLKTFRNGAFLVDKKQETSIPDVYAIGDCATVYDNAINDTNYIALASNALRSGIVAGHNAAGHKLESLGVQGSNGISIFGLNMVSTGLTQEKAKRFGYNPEVTAFTDFQKASFIEHDNYPVTLKIVYDKDSRLVLGAQMASKEDMSMGIHMFSLAIQEKVTIERLALLDYFFLPHFNQPYNYMIKAALKAKGGSAVSKGEELFTGVVPILVELDGDVNGHKFSVSGEGEGDATYGKLTLKFICTTGKLPVPWPTLVTTLTYGVQCFSRYPDHMKQHDFFKSAMPEGYVQERTIFFKDDGNYKTRAEVKFEGDTLVNRIELKGIDFKEDGNILGHKLEYNYNSHNVYIMADKQKNGIKVNFKIRHNIEDGSVQLADHYQQNTPIGDGPVLLPDNHYLSTQSKLSKDPNEKRDHMVLLEFVTAAGITLGMDELYK*

***Nucleotide and translated amino acid sequences of positive control expression cassettes used for localisation experiments***

Full-length sequences of the positive control expression cassettes used for transient expression in protoplasts (Seq ID 20 – 23). Expression of both untargeted and translationally-fused GFP in the absence of *Sm*NOX is driven by the CaMV35S promoter and contain the *A. tumefaciens* NOSt terminator.

**SEQ ID 20** – *Nucleotide sequence of the positive control expression cassette of untargeted GFP (Free GFP; cytosolic localisation)*

tcaacatggtggagcacgacactctggtctactccaaaaatgtcaaagatacagtctcagaagatcaaagggctattgagacttttcaacaaaggataatttcgggaaacctcctcggattccattgcccagctatctgtcacttcatcgaaaggacagtagaaaaggaaggtggctcctacaaatgccatcattgcgataaaggaaaggctatcattcaagatctctctgccgacagtggtcccaaagatggacccccacccacgaggagcatcgtggaaaaagaagaggttccaaccacgtctacaaagcaagtggattgatgtgacatctccactgacgtaagggatgacgcacaatcccactatccttcgcaagacccttcctctatataaggaagttcatttcatttggagaggacacgca**atg**gtgagcaagggcgaggagctgttcaccggggtggtgcccatcctggtcgagctggacggcgacgtaaacggccacaagttcagcgtgtccggcgagggcgagggcgatgccacctacggcaagctgaccctgaagttcatctgcaccaccggcaagctgcccgtgccctggcccaccctcgtgaccaccctgacctacggcgtgcagtgcttcagccgctaccccgaccacatgaagcagcacgacttcttcaagtccgccatgcccgaaggctacgtccaggagcgcaccatcttcttcaaggacgacggcaactacaagacccgcgccgaggtgaagttcgagggcgacaccctggtgaaccgcatcgagctgaagggcatcgacttcaaggaggacggcaacatcctggggcacaagctggagtacaactacaacagccacaacgtctatatcatggccgacaagcagaagaacggcatcaaggtgaacttcaagatccgccacaacatcgaggacggcagcgtgcagctcgccgaccactaccagcagaacacccccatcggcgacggccccgtgctgctgcccgacaaccactacctgagtactcagtccaagctgagcaaagaccccaacgagaagcgcgatcacatggtcctgctggagttcgtgaccgccgccgggatcactctcggcatggacgagctgtacaaatgagcttgtcaagcagatcgttcaaacatttggcaataaagtttcttaagattgaatcctgttgccggtcttgcgatgattatcatataatttctgttgaattacgttaagcatgtaataattaacatgtaatgcatgacgttatttatgagatgggtttttatgattagagtcccgcaattatacatttaatacgcgatagaaaacaaaatatagcgcgcaaactaggataaattatcgcgcgcggtgtcatctatgttactagatcga

**SEQ ID 21** – *Nucleotide sequence of the positive control expression cassette of GFP translationally fused at the N-termini to the full-length A. thaliana rubisco small subunit gene (RbcS-GFP; chloroplast stromal localisation)*

tcaacatggtggagcacgacactctggtctactccaaaaatgtcaaagatacagtctcagaagatcaaagggctattgagacttttcaacaaaggataatttcgggaaacctcctcggattccattgcccagctatctgtcacttcatcgaaaggacagtagaaaaggaaggtggctcctacaaatgccatcattgcgataaaggaaaggctatcattcaagatctctctgccgacagtggtcccaaagatggacccccacccacgaggagcatcgtggaaaaagaagaggttccaaccacgtctacaaagcaagtggattgatgtgacatctccactgacgtaagggatgacgcacaatcccactatccttcgcaagacccttcctctatataaggaagttcatttcatttggagaggacacgca**atg**gcttcttctatgctgtccagcgctactatggtggcatctcccgcacaggctaccatggtcgctcccttcaatggcttgaaatcttccgctgccttccccgcaacacgcaaggcgaacaatgatatcactagcattacctctaacggaggtcgtgtcaattgtatgcaggtgtggcctcccatcggtaaaaagaagttcgaaaccctgagctacctgcctgacctgactgactctgagctggcaaaggaggttgactatctgatccgtaacaagtggattccctgcgttgagtttgagctcgagcatgggttcgtttaccgtgagcacggaaactctcccggatattacgacggtcgttactggaccatgtggaaactgccacttttcggttgcactgattctgcacaggtccttaaggaagttgaggaatgcaaaaaggagtatccaaacgcattcatccgtatcattggatttgacaacactcgccaggtgcaatgcatttcattcatcgcctacaagcccccttcgtttaccggtggatctggtggaggtagcgcggtgagcaagggcgaggagctgttcaccggggtggtgcccatcctggtcgagctggacggcgacgtaaacggccacaagttcagcgtgtccggcgagggcgagggcgatgccacctacggcaagctgaccctgaagttcatctgcaccaccggcaagctgcccgtgccctggcccaccctcgtgaccaccctgacctacggcgtgcagtgcttcagccgctaccccgaccacatgaagcagcacgacttcttcaagtccgccatgcccgaaggctacgtccaggagcgcaccatcttcttcaaggacgacggcaactacaagacccgcgccgaggtgaagttcgagggcgacaccctggtgaaccgcatcgagctgaagggcatcgacttcaaggaggacggcaacatcctggggcacaagctggagtacaactacaacagccacaacgtctatatcatggccgacaagcagaagaacggcatcaaggtgaacttcaagatccgccacaacatcgaggacggcagcgtgcagctcgccgaccactaccagcagaacacccccatcggcgacggccccgtgctgctgcccgacaaccactacctgagtactcagtccaagctgagcaaagaccccaacgagaagcgcgatcacatggtcctgctggagttcgtgaccgccgccgggatcactctcggcatggacgagctgtacaaatgagcttgtcaagcagatcgttcaaacatttggcaataaagtttcttaagattgaatcctgttgccggtcttgcgatgattatcatataatttctgttgaattacgttaagcatgtaataattaacatgtaatgcatgacgttatttatgagatgggtttttatgattagagtcccgcaattatacatttaatacgcgatagaaaacaaaatatagcgcgcaaactaggataaattatcgcgcgcggtgtcatctatgttactagatcga

**SEQ ID 22** – *Translated amino acid sequence of the positive control untargeted GFP (Free GFP; cytosolic localisation)*

**M**VSKGEELFTGVVPILVELDGDVNGHKFSVSGEGEGDATYGKLTLKFICTTGKLPVPWPTLVTTLTYGVQCFSRYPDHMKQHDFFKSAMPEGYVQERTIFFKDDGNYKTRAEVKFEGDTLVNRIELKGIDFKEDGNILGHKLEYNYNSHNVYIMADKQKNGIKVNFKIRHNIEDGSVQLADHYQQNTPIGDGPVLLPDNHYLSTQSKLSKDPNEKRDHMVLLEFVTAAGITLGMDELYK*

**SEQ ID 23** – *Translated amino acid sequence of the positive control GFP translationally fused at the N-termini to the full-length A. thaliana rubisco small subunit gene (RbcS-GFP; chloroplast stromal localisation)*

**M**ASSMLSSATMVASPAQATMVAPFNGLKSSAAFPATRKANNDITSITSNGGRVNCMQVWPPIGKKKFETLSYLPDLTDSELAKEVDYLIRNKWIPCVEFELEHGFVYREHGNSPGYYDGRYWTMWKLPLFGCTDSAQVLKEVEECKKEYPNAFIRIIGFDNTRQVQCISFIAYKPPSFTGGSGGGSAVSKGEELFTGVVPILVELDGDVNGHKFSVSGEGEGDATYGKLTLKFICTTGKLPVPWPTLVTTLTYGVQCFSRYPDHMKQHDFFKSAMPEGYVQERTIFFKDDGNYKTRAEVKFEGDTLVNRIELKGIDFKEDGNILGHKLEYNYNSHNVYIMADKQKNGIKVNFKIRHNIEDGSVQLADHYQQNTPIGDGPVLLPDNHYLSTQSKLSKDPNEKRDHMVLLEFVTAAGITLGMDELYK*
